# Structural basis for protein glutamylation by the *Legionella* pseudokinase SidJ

**DOI:** 10.1101/2021.07.16.451782

**Authors:** Michael Adams, Rahul Sharma, Thomas Colby, Felix Weis, Ivan Matic, Sagar Bhogaraju

## Abstract

*Legionella pneumophila* (LP) avoids phagocytosis by secreting nearly 300 effector proteins into the host cytosol. SidE family of effectors (SdeA, SdeB, SdeC and SidE) employ phosphoribosyl serine ubiquitination to target an array of host Rab GTPases and innate immune factors. To suppress the deleterious toxicity of SidE enzymes in a timely manner, LP employs a metaeffector named SidJ. Upon activation by host Calmodulin (CaM), SidJ executes an ATP-dependent glutamylation to modify the catalytic residue Glu860 in the mono-ADP-ribosyl transferase (mART) domain of SdeA. SidJ is a unique glutamylase that adopts a kinase-like fold but contains two nucleotide-binding pockets. There is a lack of consensus about the substrate recognition and catalytic mechanism of SidJ. Here, we determined the cryo-EM structure of SidJ in complex with its substrate SdeA in two different states of catalysis. Our structures reveal that both phosphodiesterase (PDE) and mART domains of SdeA make extensive contacts with SidJ. In the pre-glutamylation state structure of the SidJ-SdeA complex, adenylylated E860 of SdeA is inserted into the non-canonical (migrated) nucleotide-binding pocket of SidJ. Structure-based mutational analysis indicates that SidJ employs its migrated pocket for the glutamylation of SdeA. Finally, using mass spectrometry, we identified several transient autoAMPylation sites close to both the catalytic pockets of SidJ. Our data provide unique insights into the substrate recognition and the mechanism of protein glutamylation by the pseudokinase SidJ.

## Introduction

Protein ubiquitination is a fundamental post-translational modification process that regulates a host of cellular processes including protein turnover, DNA repair, vesicular transport, innate immunity and cell cycle^1,2^. Pathogenic bacteria do not possess any ubiquitin (Ub) system of their own but often secrete effector proteins that hijack the host Ub pathways to evade host defense mechanisms and facilitate intracellular replication of bacteria^3^. LP secretes nearly 300 effector proteins into the host cytosol during infection targeting a multitude of host pathways often by employing atypical biochemical activities^4,5^. The SidE family of *Legionella* effectors carry out non-canonical ubiquitination to target multiple endoplasmic reticulum (ER) membrane resident proteins, trigger ER fragmentation, and recruit the ER-derived vesicles to the *Legionella* containing vacuole (LCV)^6–8^.

SidE enzymes catalyze Ub transfer to substrate serines without the need of cellular E1 and E2 enzymes^6^. SidEs, however, need multiple steps to catalyze this novel ubiquitination. In the first step of ubiquitination, the mART domain in SidEs ADPribosylates Arg42 of Ub using NAD^+^ as a co-factor. Subsequently, the PDE domain uses ADPribosylated Ub as a substrate and performs a histidine intermediate-driven phosphoryl transfer reaction resulting in phosphoribosyl (PR) ubiquitination of target serine residues^6–8^. SidEs, including the most studied member of the family SdeA, are also deemed toxic to eukaryotic cells due to the phosphoribosylation of Ub activity which renders the canonical host Ub system inactive^7^. A systematic study using yeast toxicity analysis has revealed that LP possesses at least 14 of the so-called metaeffectors which regulate the activity of some of legionella’s own toxic effector proteins^9^. SidJ is one such metaeffector of LP that represses the toxicity of the SidE family, and effectively rescues SidE family-induced lethality in yeast^10–12^. Interestingly, deletion of SidJ shows a more pronounced growth phenotype compared to deletion of SidEs in *Legionella* infection experiments conducted in amoeba and macrophages^10,13^. During the infection, SidEs first get secreted into the cytoplasm where they ubiquitinate several ER-resident proteins and exert their toxicity^7,14^. Levels of SidEs in the host cytoplasm peak 30 minutes post-infection and diminish thereafter in a SidJ-dependent manner^10,14^. SidJ deletion, therefore, results in a marked *Legionella* intracellular growth phenotype presumably due to persistent, uncurbed toxicity of SidEs in the host cytoplasm in the absence of SidJ^11,12,15^. Although the deubiquitination (DUB) activity of SidJ remains uncorroborated^16–18^, recent studies have shown that LP contains two effectors DupA and DupB which act as *bona fide* DUBs for PR ubiquitination^19,20^.

More recently, we and other groups have independently shown that SidJ catalyzes ATP-dependent glutamylation of SdeA catalytic residues Glu860 and Glu862 thereby inhibiting ADP-ribosylation of Ub and hence PR-ubiquitination^16–18,21^. Glutamylation activity of SidJ is strictly dependent upon the host protein Calmodulin (CaM) which binds to SidJ with nanomolar affinity. Structural studies of SidJ in complex with CaM have revealed that SidJ contains a kinase-like domain (KD) consisting of equivalent N and C-lobes sandwiching a canonical ATP-binding pocket. CaM binds to the C-terminal domain (CTD) of SidJ and stabilizes the canonical pocket allosterically. Intriguingly, SidJ was also revealed to contain a non-canonical, migrated nucleotide-binding pocket within the C-lobe of the KD. Mutagenesis of residues lining both the canonical pocket and the migrated pocket resulted in loss of SidJ’s glutamylation activity, indicating both nucleotide-binding pockets play an important role in catalysis^17,18^. It has been suggested that SidJ-mediated glutamylation occurs through a two-step mechanism: in the first step, SidJ adenylylates (attachment of AMP) SdeA forming a transient intermediate; in the second step, free L-Glutamate launches a SidJ-enabled nucleophilic attack leading to glutamylation and causing AMP release^18^. But the basis for the specificity of SidJ towards SidE enzymes and the precise roles of the two nucleotide-binding pockets in coordinating the catalysis remain unknown. Black et al., have hypothesized that the canonical pocket of SidJ is responsible for carrying out initial ATP hydrolysis coupled to the adenylylation of Glu860 of SdeA, whereas the migrated pocket is suggested to be responsible for the glutamylation reaction^18^. However, Sulpizio et al., have found that mutating residues in the migrated pocket also disrupted the adenylylation of Glu860 of SdeA and argued that the migrated pocket could be an allosteric nucleotide-binding site necessary to stabilize the canonical pocket which executes both adenylation and glutamylation reactions^17^.

Here, we trapped the pre-glutamylation reaction intermediate of SidJ/CaM-SdeA complex for structural studies by introducing a single point mutation in SidJ. We purified a stable complex of adenylylated SdeA bound to SidJ and using cryogenic electron microscopy (cryo-EM), we determined a 2.9 Å structure of this complex. The structure reveals that the adenylylated E860 residue of SdeA is inserted into the migrated pocket of SidJ and poised for glutamylation. We have also determined the post-catalytic state structure of SidJ/CaM and SdeA, in which we captured apo-SidJ bound to glutamylated SdeA likely poised for the next adenylylation and glutamyl chain extension reactions. Both PDE and mART domains of SdeA interact with SidJ, explaining why a truncation in PDE rendered SdeA resistant to SidJ in previous yeast toxicity rescue experiments^11^. Our structure-based mutational analysis revealed that SidJ, despite adopting a pseudo-kinase fold, uses its non-canonical migrated pocket for glutamylation of its substrates. Furthermore, using mass spectrometry we found that SidJ undergoes autoAMPylation on specific lysine and glutamate residues proximal to both the catalytic pockets.

## Results

### Trapping SidJ during the glutamylation of SdeA

SidJ and SdeA did not show any detectable co-elution when assayed using *in cellulo* co-IPs, indicating a transient nature of the interaction possibly limited to the moment of catalysis (Figure S1A). We hypothesized that a mutant form of SidJ might be able to alter the glutamylation reaction kinetics to make the interaction between SidJ and SdeA less transient. To test this, we mutated a number of SidJ residues both in the canonical pocket (D542A, K367A, Q350A, T353A, K370A, Y452A) (Figure S1B) and in the migrated pocket (E565A, H492A) (Figure S1C) and purified these SidJ mutants in complex with CaM. We then used these individual mutant SidJ-CaM complexes in an *in vitro* reaction with SdeA (residues 231 to 1190 containing both PDE and mART domains) and ATP and subjected the reaction mixture to analytical size-exclusion chromatography (Figure S1D). As expected, the WT SidJ-CaM complex and SdeA did not form a complex under these conditions and eluted as heterodimer and monomer respectively (Figure S1D-E). Among the SidJ mutants tested, a mutant of the migrated pocket, SidJ E565A-CaM incubated with SdeA and ATP eluted in two separate but overlapping peaks (Figure 1A). The elution peak corresponding to the estimated molecular weight of ~200kDa contained stoichiometric amounts of SdeA, SidJ and CaM (Figure 1A) indicating the formation of a ternary complex. Notably, SidJ E565A-CaM and SdeA did not form the ternary complex under similar conditions when L-glutamate was added to the reaction mix or when we replaced ATP in the reaction mix with the non-cleavable ATP analogue ApCpp (Figure 1B).

**Figure 1.**
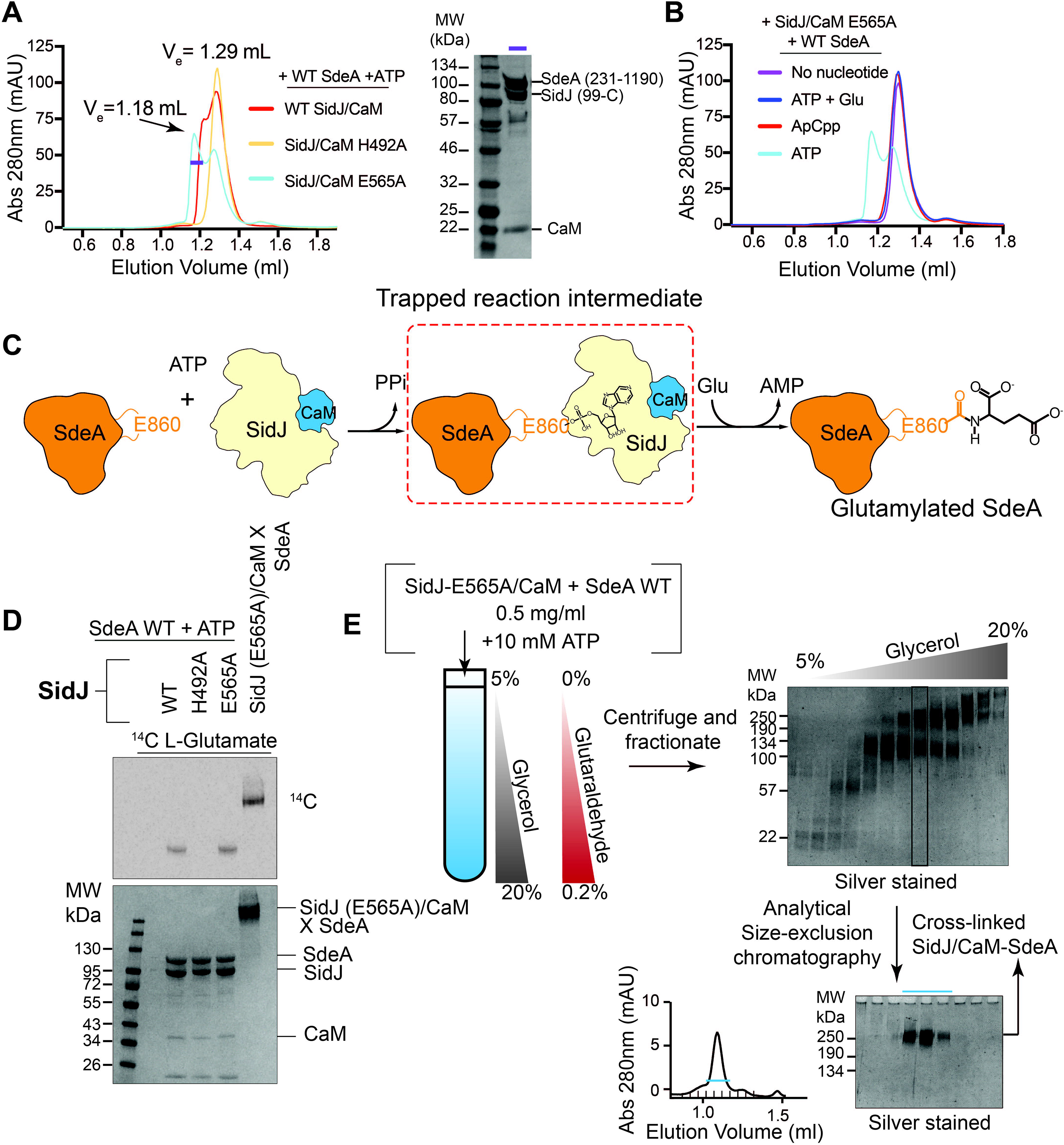
SidJ-E565A/CaM and SdeA form a stable reaction intermediate complex A) Size-exclusion chromatography (SEC) profiles of SidJ/CaM and SdeA in the presence of ATP. The highlighted fraction in the SidJ/CaM E565A + SdeA sample is shown on SDS-PAGE (right) B) SEC profiles of SidJ/CaM E565A + SdeA in the presence of various cofactors C) Schematic representation of the reaction scheme of SidJ-mediate glutamylation of SdeA highlighting the trapped adenylylated reaction intermediate between SidJ/CaM and SdeA D) Incorporation of [^14^C]-Glu into SdeA catalyzed by SidJ using SidJ WT, SidJ H492A, SidJ E565A, and cross-linked SidJE565A/CaM/SdeA, with SdeA tethered to the migrated pocket (Prepared as in 1E). Samples were separated by SDS-PAGE and visualized by Coomassie stain (Bottom) or autoradiography (Top) E) Flow chart of the sample preparation process of SidJ-E565A/CaM + SdeA for Cryo-EM. From left to right, samples are incubated and then cross-linked via GraFix method. Fractions are analyzed via SDS-PAGE, and further purified via SEC

In WT SidJ, hydrolysis of ATP leads to adenylylation of the target glutamate of SdeA leading to the formation of an acyl-AMP moiety (Figure 1C). Subsequently, the amino group of a free L-Glutamic acid molecule launches a SidJ-enabled nucleophilic attack on the acyl-AMP intermediate, resulting in glutamylation of the target residue via an isopeptide bond. Our data (Figure 1A-B) indicates that SidJ E565A-CaM-SdeA heterotrimeric complex is likely a reaction intermediate that occurs after ATP hydrolysis but before glutamylation by SidJ. Surprisingly, SidJ E565A glutamylates SdeA to the same extent as WT in our *in vitro* glutamylation assays (Figure 1D) but in the absence of kinetic measurements we cannot rule out an altered glutamylation rate for SidJ E565A mutant. To study this pre-glutamylation complex using cryo-EM, we applied gradient fixation (GraFix)^22^ on the SidJ E565A-CaM-SdeA complex and purified the cross-linked complex further using analytical size-exclusion chromatography (Figure 1E). The purified SidJ E565A-CaM-SdeA complex elutes in a single size-exclusion peak and runs as one cross-linked species in SDS-PAGE (Figure 1E). Importantly, the cross-linked SidJ E565A-CaM-SdeA complex is still active as assayed through *in vitro* glutamylation reactions (Figure 1D) indicating that GraFix does not completely incapacitate the complex.

### Overall structure of SidJ-CaM-SdeA ternary complex in pre-glutamylation state

The purified SidJ E565A/CaM-SdeA complex was analyzed by single particle cryo-EM yielding a density map with a nominal resolution of 2.9 Å (Figure S2, Table S1). The obtained map was used to rigid body fit both SidJ/CaM (PDB:6oqq)^16–18,21^ and SdeA 231-1190 (PDB:5yij)^23–26^ and refined. The C-terminal region of SdeA corresponding to residues 905 to 1190 is not resolved in the EM map (Figure S2) and the alpha-helical lobe (AHL corresponding to residues 591-757) of the mART domain is also not adequately resolved in the map. The refined structure contains one molecule each of SdeA, SidJ and CaM with the adenylylated E860 of SdeA pointing directly into the migrated pocket of SidJ (Figure 2A-C). The isolated SidJ/CaM structure superimposes well with the SidJ/CaM in complex with SdeA with a mean r.m.s.d of 0.5 Å over 744 C-α atoms (Figure S3A). The SdeA structure that was previously resolved through X-ray crystallography also superimposes well with SdeA in complex with SidJ with a mean r.m.s.d of 1.3 Å over 665 C-α atoms (Figure S3B). The overall structure of SidJ/CaM-SdeA heterotrimer resembles a triangular pyramid with the apex formed of the SdeA PDE domain and the base of the pyramid formed by the “back-face” of SidJ/CaM (Figure 2A-B). The interface of SidJ and SdeA contains both hydrophobic and polar interactions with a combined buried surface area of ~2100 Å^2^. CaM in the SidJ/CaM-SdeA complex adopts a similar conformation as seen in SidJ/CaM complex alone and is not involved in any direct contact with SdeA. The interaction between SidJ and SdeA is mediated through multiple domains of both SidJ and SdeA. The PDE domain and mART domain of SdeA interact with the C-lobe of SidJ KD (Figure 2B). Intriguingly, the N-lobe of SidJ KD and the canonical pocket of SidJ, which is implicated in the catalysis, do not make any contacts with SdeA (Figure 2B). This is possibly because the trapped SidJ/CaM-SdeA complex represents a state of catalysis after the adenylylation of SdeA and before the glutamylation which was hypothesized to occur at the migrated pocket^18^. Although glutamylation target residue E860 of SdeA is clearly seen adenylylated and inserted into the migrated pocket of SidJ, there is no discernible density for adjacent residues 854-859 of the mART catalytic loop of SdeA (Figure 2C). This implies that the specificity of SidJ towards E860 of SdeA is achieved through other sites of SidJ-SdeA interactions distant from the catalytic site and likely independent of the sequence immediately surrounding the target glutamate. Accordingly, an SdeA ΔPDE construct (spanning residues 531-1190) containing the intact mART domain could only be glutamylated negligibly by SidJ, indicating that SidJ relies on interactions with both PDE and mART domains of SdeA to specifically glutamylate E860 of SdeA (Figure 2D). Residue H492 in SidJ which coordinates the α-phosphate group of AMP in the migrated pocket (Figure S1C) was shown in previous studies to be essential for SidJ catalysis^17,18^, hence we used SidJ H492A as a negative control in our biochemical assays. Consistent with the involvement of the SdeA PDE domain in interaction with SidJ, previous yeast toxicity experiments have also observed that SidJ could not rescue the toxicity of SdeA construct lacking part of the PDE domain^11^.

**Figure 2.**
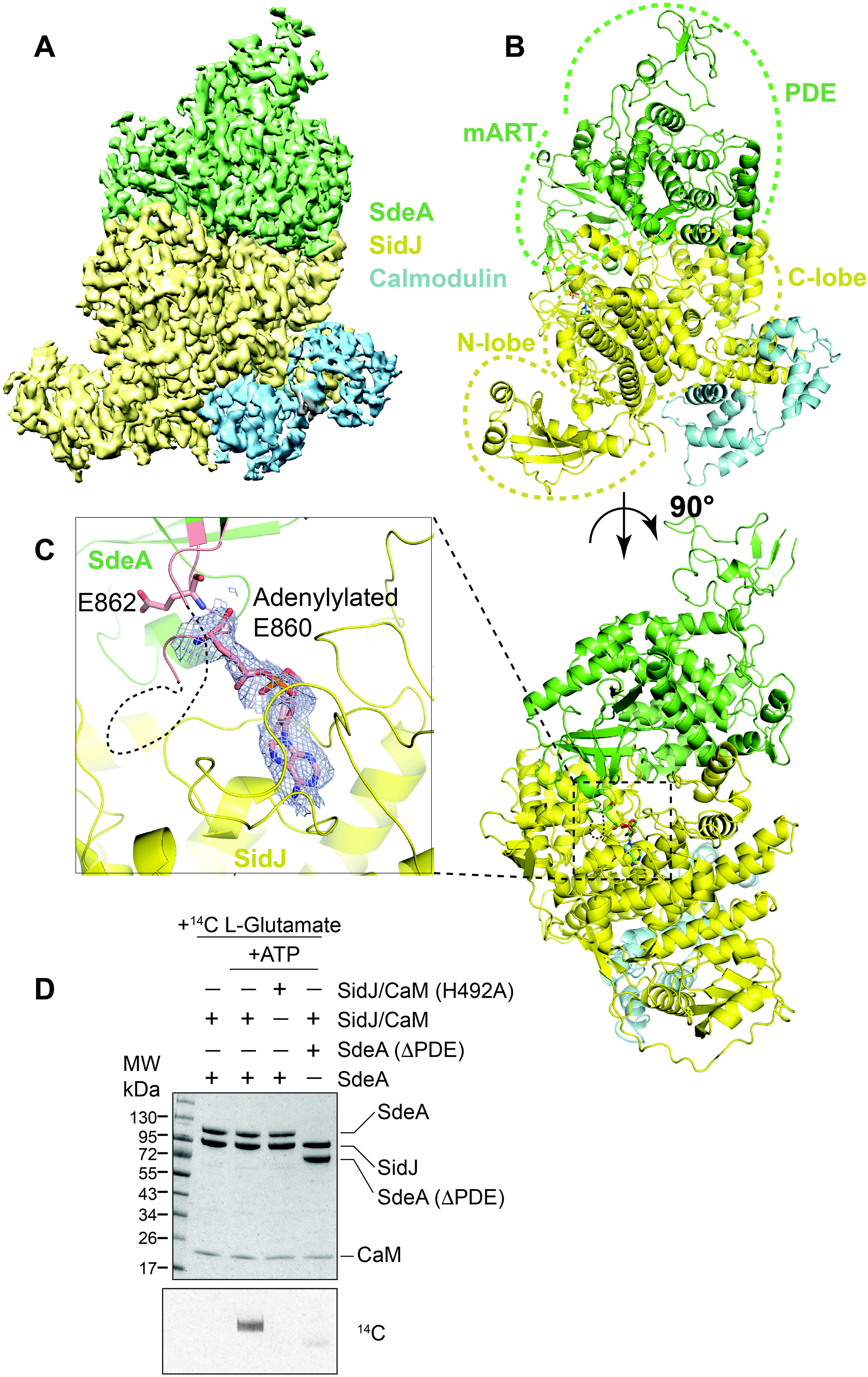
Adenylylated SdeA E860 binds to the migrated pocket of SidJ A) Cryo-EM map of SidJ-E565A (Yellow), Calmodulin (Teal), and SdeA (Green) B) Overall structure of the reaction intermediate SidJ-E565A/CaM/SdeA heterotrimer. Both N- and C-lobe of SidJ (Yellow), as well as mART and PDE lobes of SdeA (Green) are highlighted, with Calmodulin (Teal) bound to the C-terminus of SidJ C) Closer view of the migrated nucleotide-binding pocket of SidJ (Yellow), and Adenylylated SdeA E860 (Stick representation, Green) inserted. The cryo-EM density of E860 of SdeA and AMP is shown in mesh. SdeA catalytic loop is colored in salmon. E862, another glutamylation target of SidJ is also shown. D) Incorporation of [14C]-Glu into SdeA with and without its PDE domain present. Reaction components were separated by SDS-PAGE and either visualized by Coomassie stain (Top) or autoradiography (Bottom)

### Characterization of SidJ-SdeA interface

The interface of SidJ and SdeA can be divided into three major contact sites spanning multiple domains of both proteins (Figure 3A). Site-1 involves an insertion loop in SidJ spanning residues 290 to 304 that protrudes outwards from the surface of SidJ and inserts into the groove between the PDE and the mART domains of SdeA (Figure 3B). Interestingly, a previous yeast screening experiment has found that mutation of P290, which is involved in the formation of a sharp kink at the start of this insertion loop, renders SidJ deficient in rescuing the toxicity of SdeA^12^ (Figure 3B). SidJ R293 at the tip of this insertion loop participates in a hydrogen bonding network involving SdeA Q572 and the backbone carbonyl of SdeA Y235 in its PDE domain (Figure 3C). Mutating R293 of SidJ did not affect the glutamylation of SdeA as assayed through *in vitro* glutamylation assays using C^14^ labeled L-glutamate (Figure 3D, right). Deletion of the extending insertion loop in SidJ (SidJ Δ291-300) resulted in only a minimal reduction of glutamylation *in vitro*, indicating that this loop does not play a critical role in SidJ-SdeA interaction and other interaction sites may complement in its absence (Figure 3D, left). Site-1 of SidJ-SdeA interaction also involves F566 of SdeA making hydrophobic contacts with M696 and Y699 of SidJ (Figure 3C). Additionally, T236 of SdeA is within hydrogen bonding distance to the backbone carbonyl of E294 of SidJ (Figure 3C). Mutating T236 or F566 of SdeA resulted in a strong reduction of SidJ-mediated glutamylation *in vitro* (Figure 3D, right).

**Figure 3.**
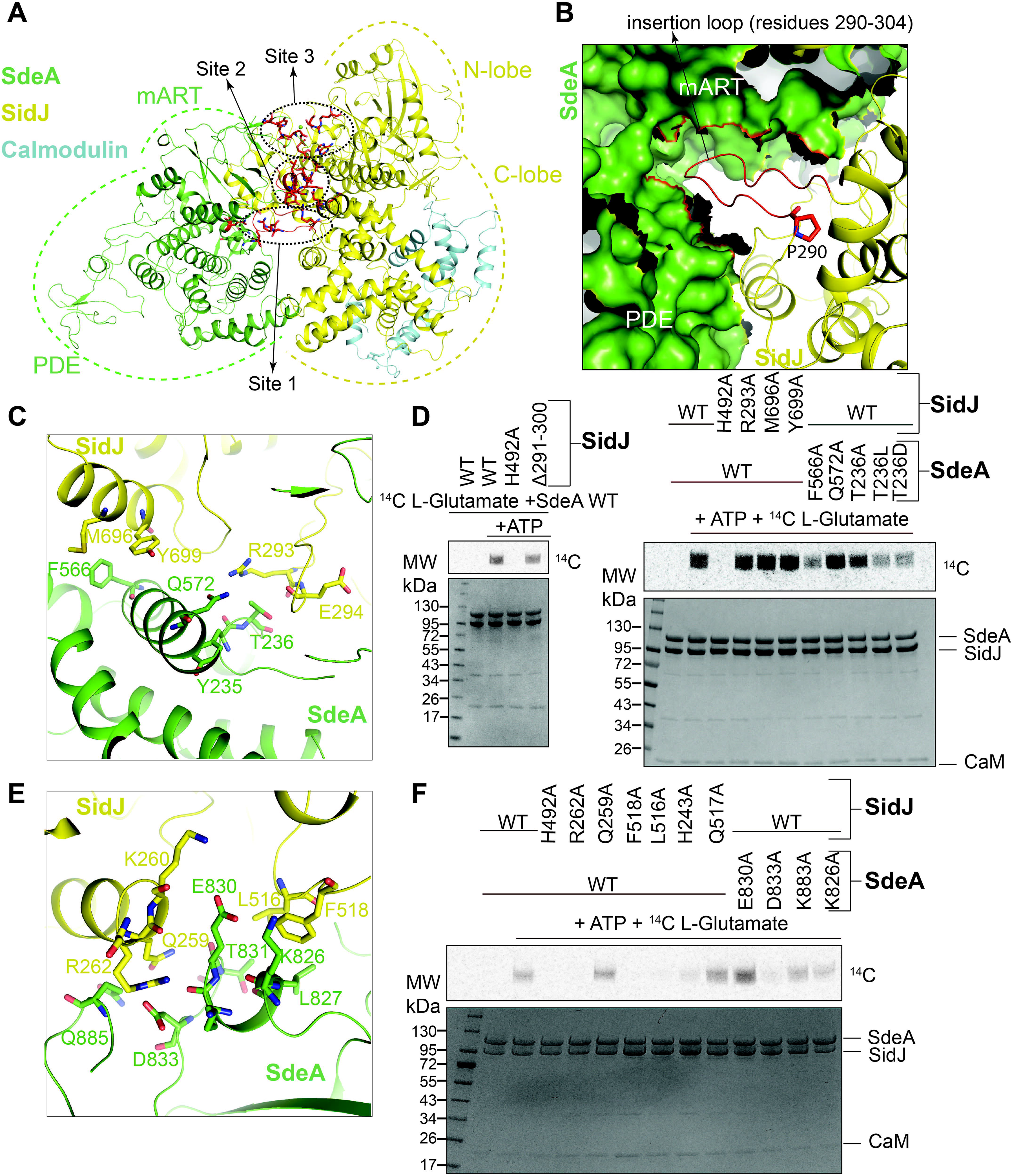
SidJ and SdeA form an extensive binding interface A) Overview of SidJ/CaM and SdeA intermediate complex showing the three binding sites between SidJ and SdeA. Site 1 being the loop insertion site (Shown in 3B,C), site 2 being the mART site (Shown in 3E), and site 3 being the migrated pocket site (Shown in 4A) B) The insertion loop of site 1 (Red) of SidJ inserting itself into the cleft between SdeA’s PDE and mART lobes (Shown in green, surface representation). SidJ Proline 290 is shown in stick representation. C) Detailed view of the SidJ-SdeA Interaction site 1-the insertion loop of SidJ (Yellow) and SdeA (green) with key residues shown D) Incorporation of [^14^C]-Glu into SdeA catalyzed by SidJ, with and without the insertion loop (Left) and using mutations on both SidJ and SdeA from site 1 (Right). Samples were separated by SDS-PAGE and visualized by Coomassie stain (Bottom) or autoradiography (Top) E) Detailed view of the SidJ-SdeA Interaction site 2-the interactions between SidJ C-lobe (Yellow) and SdeA mART, with key residues shown F) Incorporation of [^14^C]-Glu into SdeA catalyzed by SidJ using mutations on both SidJ and SdeA from site 2. Samples were separated by SDS-PAGE and visualized by Coomassie stain (Bottom) or autoradiography (Top)

Site-2 of interaction between SidJ and SdeA mainly involves a short helical insertion in the mART domain of SdeA spanning residues 825-833 mediating interactions with the C-lobe of SidJ KD (Figure 3A,E). Specifically, these interactions include two salt bridges between R262, K260 of SidJ and D833, E830 of SdeA respectively. Site-2 also includes a tight hydrogen bonding network with the side chains of residues SidJ Q259, SdeA Q885 and the backbone carbonyl of SdeA T831. In addition, residues SidJ F518, L516 pack against SdeA L827 and the aliphatic part of SdeA K826 constituting a hydrophobic interface between SidJ and SdeA. Single point mutations of a few residues involved in Site-2 resulted in loss of glutamylation activity *in vitro* (Figure 3F).

The migrated pocket of SidJ is bound tightly to the adenylylated E860 of SdeA and forms site-3 of interaction between SidJ and SdeA (Figure 4A). AMP assumes a similar conformation in SidJ’s migrated pocket as it does in the previously described crystal structures of SidJ/CaM alone^18^ (Figure S4A). Surprisingly, there are only a few contacts between SidJ and SdeA in site-3 apart from the ones involving the AMP moiety covalently linked to SdeA. A key hydrogen bonding network involving Q851 of SdeA, the backbone carbonyl of E860 of SdeA and Y732, N733 of SidJ positions the glutamylation target residue SdeA-E860 in the migrated pocket of SidJ (Figure 4A). Mutating residues involved in this hydrogen bonding network compromises SdeA glutamylation by SidJ (Figure 4B). Mg^2+^ ion and the side chain of SidJ R500 engage in a tight coordination with the α-phosphate of the AMP, likely increasing the reactivity of the acyl-AMP intermediate towards the amino-group of the incoming glutamate. Surprisingly, the introduced SidJ mutation E565A in the SidJ/CaM-SdeA complex did not cause any noticeable changes in the migrated pocket of SidJ compared to previously resolved SidJ structures (Figure S4B). Although we are not able to pinpoint why SidJ E565A binds SdeA less transiently in this reaction intermediate compared to SidJ WT, we speculate that the altered microenviroment of SidJ’s migrated pocket in E565A increases the residence time of adenylylated SdeA on SidJ (Figure S4C).

**Figure 4.**
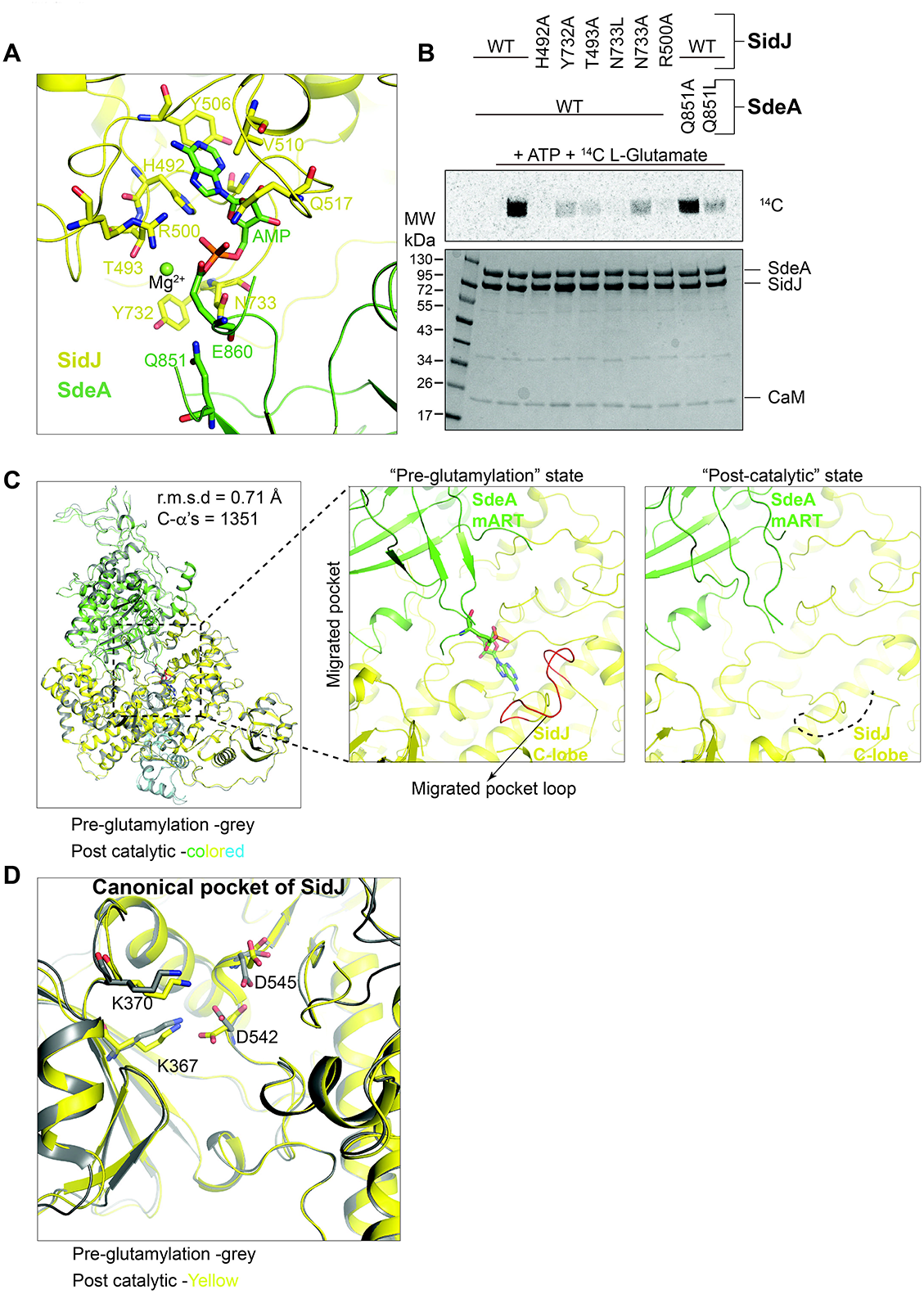
SidJ catalyzes glutamylation in the migrated pocket A) Detailed view of the SidJ-SdeA Interaction site 3-Adenylylated E860 of SdeA (Green) inserted into the migrated pocket of SidJ (Yellow), with key residues shown B) Incorporation of [^14^C]-Glu into SdeA catalyzed by SidJ using mutations on both SidJ and SdeA from site 3. Samples were separated by SDS-PAGE and visualized by Coomassie stain (Bottom) or autoradiography (Top) C) Structural comparison between SidJ-E565A (Yellow) / CaM + SdeA (Green) in its pre-glutamylation and post-catalytic states. Highlighted is a loop of SidJ (Red) located in the migrated pocket that undergoes structural change post-catalysis D) Overlay of SidJ canonical pocket in pre-glutamylation (Grey) and post-catalytic (Yellow) state, with key residues highlighted

### Structural evidence for SdeA glutamylation at the migrated pocket of SidJ

The SidJ/CaM-SdeA pre-glutamylation state structure described above strongly supports the hypothesis that SidJ’s migrated pocket is the site of SdeA E860 glutamylation. We further probed the catalytic role of the migrated pocket by exploiting the GraFixed SidJ/CaM-SdeA complex sample which still performs glutamylation reaction (Figure 1D). We added L-glutamate to the prepared SidJ/CaM-SdeA pre-glutamylation complex right before applying the sample to the EM grid and analyzing it by single particle cryo-EM (Figure S5, Table S1). Interestingly, after 3D classification of the particles, we observed two well resolved particle classes, Class I (15% of total particles) contained both SidJ/CaM and SdeA well resolved and Class II (17% of total particles) represented only SidJ/CaM with discernible density while SdeA was found to be disordered, presumably dissociated from SidJ but still tethered to it due to GraFix (Figure S5). We pursued the processing of Class I particles to obtain a 3.7Å reconstruction of SidJ/CaM-SdeA in a post-catalytic (PC) state (Figure S5). Overall, the PC state structure of SidJ/CaM-SdeA superposes well with the pre-glutamylation state structure with an r.m.s.d of 0.71Å over 1351 C-α atoms with noticeable differences only in the migrated pocket region of SidJ and the mART domain of SdeA (Figure 4C). Interestingly, post-glutamylation, the SidJ loop (residues 492-501) making up the migrated pocket becomes disordered in the structure, while the canonical pocket architecture remains intact (Figure 4C-D). Consistent with the idea that AMP releases upon SdeA glutamylation^18^, the electron density corresponding to AMP in SidJ’s migrated pocket is missing in the post-catalytic state structure of SidJ/CaM-SdeA (Figure 4C). There is, however, no electron density in the post-catalytic SidJ/CaM-SdeA structure for the added L-glutamate near SdeA E860, likely because of the flexible nature of the modification.

### Characterization of autoAMPylation of SidJ

Having gained structural insights into the second step of SidJ-mediated SdeA glutamylation, we sought to understand how the first step (SdeA adenylylation) of SidJ catalysis occurs. Previous studies have proposed that SdeA adenylylation is catalyzed by the canonical pocket of SidJ^17,18^. Interestingly, Sulpizio et al., have reported that SidJ possesses autoAMPylation activity which is chemically identical to SdeA adenylylation^17^. It was also shown that mutating both canonical pocket (D542) and migrated pocket (H492) residues renders SidJ deficient in this autoAMPylation activity^17^. We probed the nature of autoAMPylation of SidJ and its relevance in SidJ catalysis. We first tested the stability of the autoAMPylated SidJ and found that it is both heat and acid labile (Figure 5A). This indicates that the AMP is attached to SidJ through an unstable bond and autoAMPylated SidJ is transient in nature. Interestingly, compared to SidJ WT, we noticed that a migrated pocket mutant SidJ R500A exhibits marked increase in SidJ autoAMPylation and also acyl adenylylate formation with SdeA (Figure S6). Using both SidJ WT and R500A proteins, we then aimed to characterize the nature of autoAMPylation sites in SidJ using mass spectrometry. We reacted SidJ (WT or R500A) with ATP and subjected the proteins to tryptic digestion followed by LC-MS/MS analysis. Surprisingly, several transient autoAMPylation sites on residues close to the two catalytic pockets of SidJ were detected in both WT and R500A proteins (Figure 5B-C, S7). In agreement with the radioactive assays (Figure S6), we obtained more spectra for AMPylated peptides in SidJ R500A compared to WT SidJ (Figure 5C-D, S7). Yet, we found two peptides of SidJ that are AMPylated in both WT and the R500A mutant (Figure 5B-C, S7). One of these peptides (368-VQKRGEPK-375) lines the canonical pocket of SidJ (Figure 5B,D) and we could precisely localize the site of AMPylation on this peptide to the residue K370 using ETD (Electron Transfer Dissociation) fragmentation (Figure 5B, S7A). AutoAMPylation of SidJ was also detected in a peptide (480-EGIMFPQLADIFHTHFGEDEREDK-503 (S7B-C, S8A) forming a long loop (referred as bridging peptide from now on) that originates at the canonical pocket and extends into a crucial part of the migrated pocket (Figure 5C). We obtained several HCD (Higher-energy collisional dissociation) fragmentation spectra of the AMPylated bridging peptide in SidJ R500A mutant (S7B-C). Although we could not precisely localize the modification to a specific residue in the bridging peptide (because HCD fragmentation breaks off AMP from the modified residue, see Methods), we could narrow down the site through ETD fragmentation to a three residue stretch 497-EDE-499 close to the migrated pocket of SidJ (S8A). In reactions containing both SidJ and SdeA, we could also obtain an HCD spectrum showing the AMPylation of the catalytic peptide in SdeA with likely target residue as E862 (Figure S8B) providing the strongest evidence yet that SdeA is adenylylated in the course of glutamylation.

**Figure 5.**
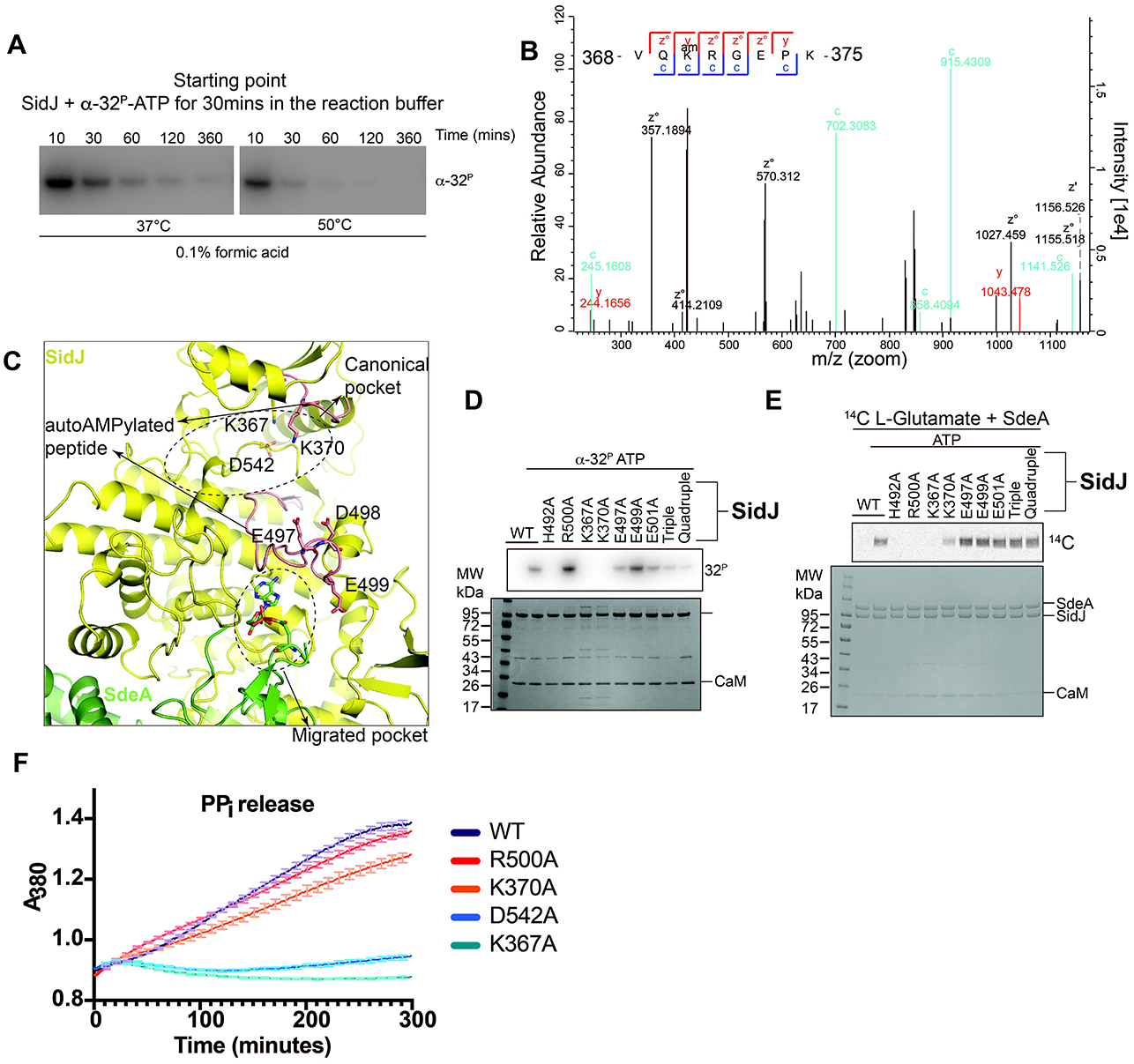
Characterization of SidJ autoAMPylation A) Time-course of SidJ auto-AMPylation in an acidic environment at different temperatures. Reactions were performed using SidJ WT + CaM and α-[32P]-ATP. Samples were separated by SDS-PAGE and visualized by autoradiography B) ETD fragmentation spectrum of SidJ peptide VQKRGEPK, with the AMPylation site K370 highlighted. AutoAMPylation reactions of SidJ WT were analyzed using LC-MS/MS. C) Detailed view of SidJ auto-AMPylation sites, with peptides identified in mass spectrometry shown in red, and modification sites highlighted D) AutoAMPylation of SidJs. CaM+SidJ WT or indicated mutants were reacted with α-[32P]-ATP. Samples were separated by SDS-PAGE and visualized by Coomassie stain (Bottom) and autoradiography (Top) E) Incorporation of [14C]-Glu into SdeA catalyzed by SidJ using SidJ WT and mutants of the auto-AMPylation sites. Samples were separated by SDS-PAGE and visualized by Coomassie stain (Bottom) and autoradiography (Top) F) Pyrophosphate release assay with SidJ and mutants indicated. The assay was performed using the EnzCheck™ pyrophosphate assay kit (ThermoFischer Scientific). The kit components (see methods) react with the PPi in solution and release ribose 1-phosphate and 2-amino-6-mercapto-7-methylpurine which shows absorption peak at 380nm.

We then performed site-directed mutagenesis on SidJ autoAMPylation sites to probe if they affect autoAMPylation levels of SidJ and glutamylation of SdeA. Since there is a cluster of glutamates (E497, E499, E501 and E565) concentrated in SidJ in and around the bridging peptide (Figure S9), we decided to mutagenize all these glutamates alone and combined. Mutating individual glutamates in the bridging loop did not result in the decrease of autoAMPylation but mutating the cluster of glutamates resulted in a marked reduction in SidJ autoAMPylation (Figure 5D). Interestingly, mutating the canonical pocket autoAMPylation residue K370 in SidJ completely abolishes autoAMPylation, indicating that the AMPylation of K370 precedes that of the bridging peptide residues (Figure 5D). Surprisingly, none of the bridging peptide mutations in SidJ affected the glutamylation of SdeA but mutating SidJ K370 resulted in a dramatic reduction of glutamylation activity (Figure 5E). We next checked if the effect of SidJ K370A mutation on autoAMPylation and glutamylation activities is due to defective ATP hydrolysis by SidJ. The ATP hydrolysis activity of SidJ K370A is comparable to that of WT as revealed in our *in vitro* pyrophosphate release assays (Figure 5F). Mutating catalytic Lys367 and Glu542 in the canonical pocket of SidJ completely abolishes the ATP hydrolysis activity (Figure 5F) indicating a clear role of canonical pocket in ATP hydrolysis by SidJ. Based on our data, autoAMPylation of glutamates in the bridging peptide of SidJ seem to be by-products of SidJ reaction although other roles such as self-regulation cannot be excluded. Importantly, we show that the autoAMPylation target residue SidJ K370 is important for SdeA glutamylation. However, a direct correlation between autoAMPylation of SidJ on the residue K370 and SdeA glutamylation is still missing. Further experimentation into autoAMPylation of SidJ is necessary in order to understand its possible relevance in catalysis or autoregulation.

## Discussion

Here, we determined the cryo-EM structure of SidJ bound to SdeA trapped at an intermediate step of catalysis. The C-lobe of the kinase-like domain of SidJ makes extensive contacts with both the mART and the PDE domains of SdeA. Interestingly, the ubiquitin-binding surface on the SdeA mART domain overlaps with the SidJ-binding surface, indicating that ubiquitin and SidJ compete for the same surface for binding to SdeA^25^ (Figure S10). The pre-glutamylation structure has no nucleotide bound in the canonical pocket of SidJ but shows the adenylylated E860 of the SdeA mART domain inserting itself into the migrated pocket of SidJ primed for glutamylation. In the post-catalytic structure of SidJ/CaM-SdeA complex where one cycle of glutamylation occurred, SidJ exists in an apoenzyme form with no nucleotide bound in either canonical or migrated pockets (Figure 4). Importantly, part of the loop building the migrated pocket (residues 492-501) becomes disordered after a cycle of glutamylation (Figure 4C). This observation argues against the proposed allosteric role for the migrated pocket^17^ because the canonical pocket is fully formed even though the migrated pocket is not bound to any nucleotide and disordered in the post-catalytic SidJ/CaM-SdeA structure. Cryo-EM structures presented in this study provide a basis for substrate recognition of SidJ and show that the migrated pocket of SidJ is a dynamic catalytic center that binds to the reaction intermediate-adenylylated SdeA and performs glutamylation.

SidJ autoAMPylation occurs on a lysine in the canonical pocket (K370) and also likely on several glutamates in the bridging loop that connects the canonical pocket and the migrated pocket (Figure 5, S7, S8). Mutation of modified glutamate residues in SidJ reduces autoAMPylation but does not affect glutamylation activity, indicating that these autoAMPylation events are possibly either side reactions of SidJ catalysis or have some unknown role in self-regulation which is yet to be explored. Mutation of K370 completely abolishes the autoAMPylation of SidJ indicating that modification of K370 precedes that of glutamates in the bridging peptide of SidJ. SidJ K370A mutant protein shows strongly reduced glutamylation activity compared to WT SidJ (Figure 5E). We also showed that the reduced glutamylation activity of SidJ K370A mutant is not due to a defect in ATP hydrolysis (Figure 5F). However, SidJ K370A mutant protein still showed residual glutamylation activity (Figure 5E), so autoAMPylaiton of K370 is not essential for SidJ-mediated glutamylation of SdeA. It is also puzzling that SidJ migrated pocket mutant H492A shows complete lack of autoAMPylation (Figure 5D). This indicates that the migrated pocket of SidJ may also play a role in autoAMPylation in addition to catalyzing the glutamylation step. Further studies are necessary to understand the role of the transient autoAMPylation sites our study identified in SidJ. Interestingly, DNA ligases also carry out lysine AMPylation as an intermediate reaction before AMP is transferred to a DNA 5′ phosphate group and is eventually released upon DNA ligation^27,28^. Additionally, selenoprotein-O (SelO), a conserved pseudokinase was recently shown to AMPylate substrate proteins as well, but unlike SidJ, SelO targets its substrates on serine, threonine and tyrosine residues^29^.

SidJ R500A mutant protein exhibits more autoAMPylation and acyl adenylylate formation compared to WT SidJ (Figure S6). Interestingly, a recent pre-print by Osinski et al., shows that SidJ R500 co-ordinates free L-Glutamate along with R522 which plays a crucial part in SidJ-enabled nucleophilic attack by L-Glutamate^30^. This explains why SidJ R500A mutant shows increased autoAMPylation while being defective in glutamylation of SdeA.

SidJ belongs to a very small number of *Legionella* effectors whose deletion results in significant reduction in bacterial intracellular growth^10,13,14^. SidJ facilitates *Legionella* replication likely by curbing the excessive toxicity of SidEs which is exerted by their ubiquitination and ubiquitin modification activities^6–8,10,31^. Previous infection experiments indicate that compared to deletion of SidEs, deletion of SidJ in general carries a greater effect on the intracellular replication of *Legionella* in both murine macrophages and amoeba^10^. This could be due to two reasons that are not mutually exclusive, first being that SidJ targets other host cellular substrates during *Legionella* infection, secondly, prolonged persistence of toxic SidEs in the host cytoplasm due to lack of SidJ is more detrimental to *Legionella*’s replication than complete lack of SidEs. Future *Legionella* infection experiments complementing *ΔSidE* Legionella strain with SidE mutants resistant to SidJ but possessing wild-type activity might help delineate the pathophysiological role of SidJ beyond SidEs.

---

**Figure S1.**
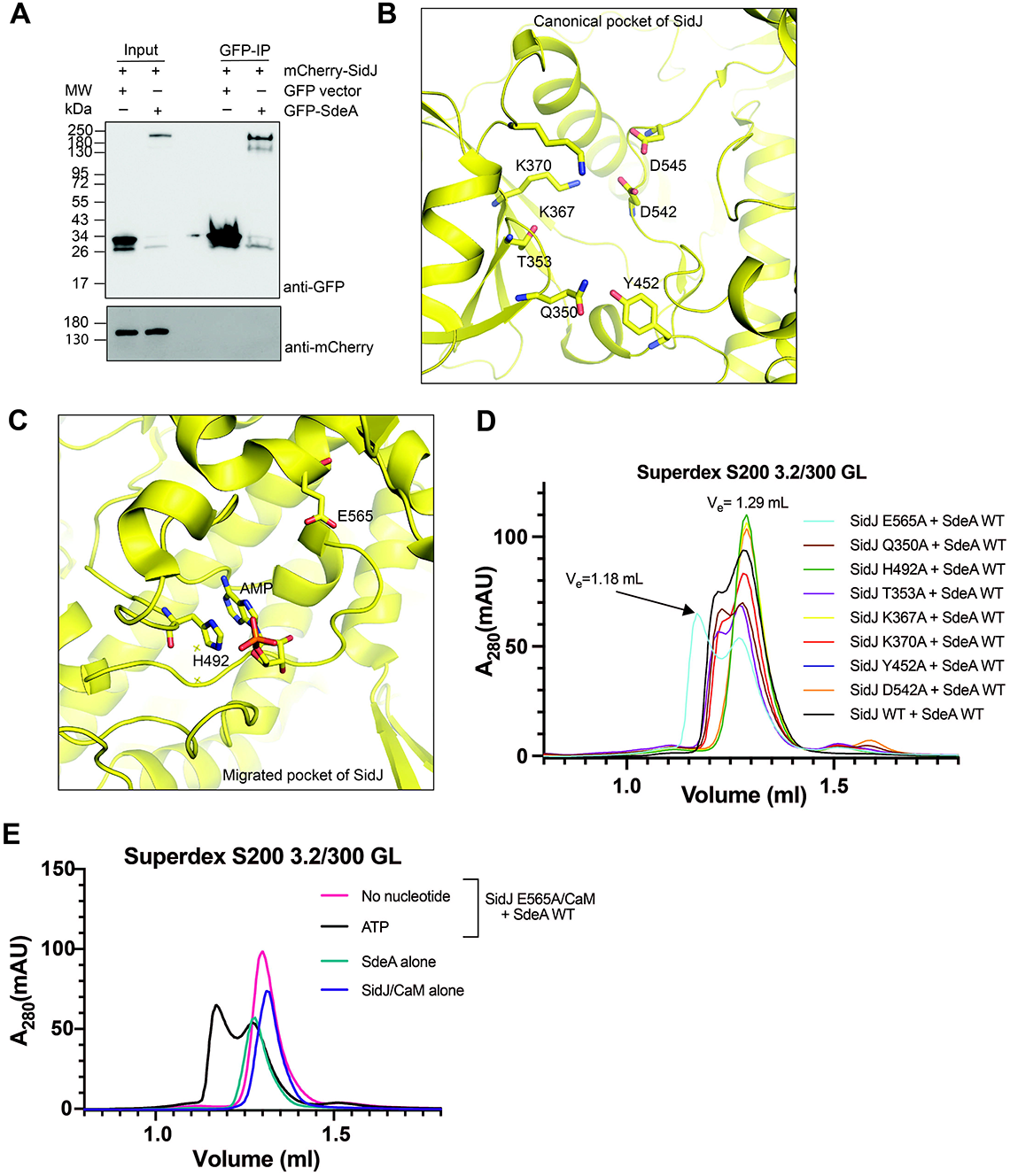
SidJ - SdeA interaction is transient A) mCherry-SidJ and either GFP-SdeA or GFP expressed in HEK293T cells. GFP IP was performed followed by blotting with anti-GFP and anti-mCherry antibodies B) Detailed view of SidJ canonical pocket with mutated residues shown C) Detailed view of SidJ migrated pocket with mutated residues shown D) SEC profiles of SidJ/CaM point mutants mixed with SdeA in the presence of ATP E) SEC profiles of SidJ-E565A/CaM and SdeA with and without ATP, as well as SidJ/CaM and SdeA alone

**Figure S2.**
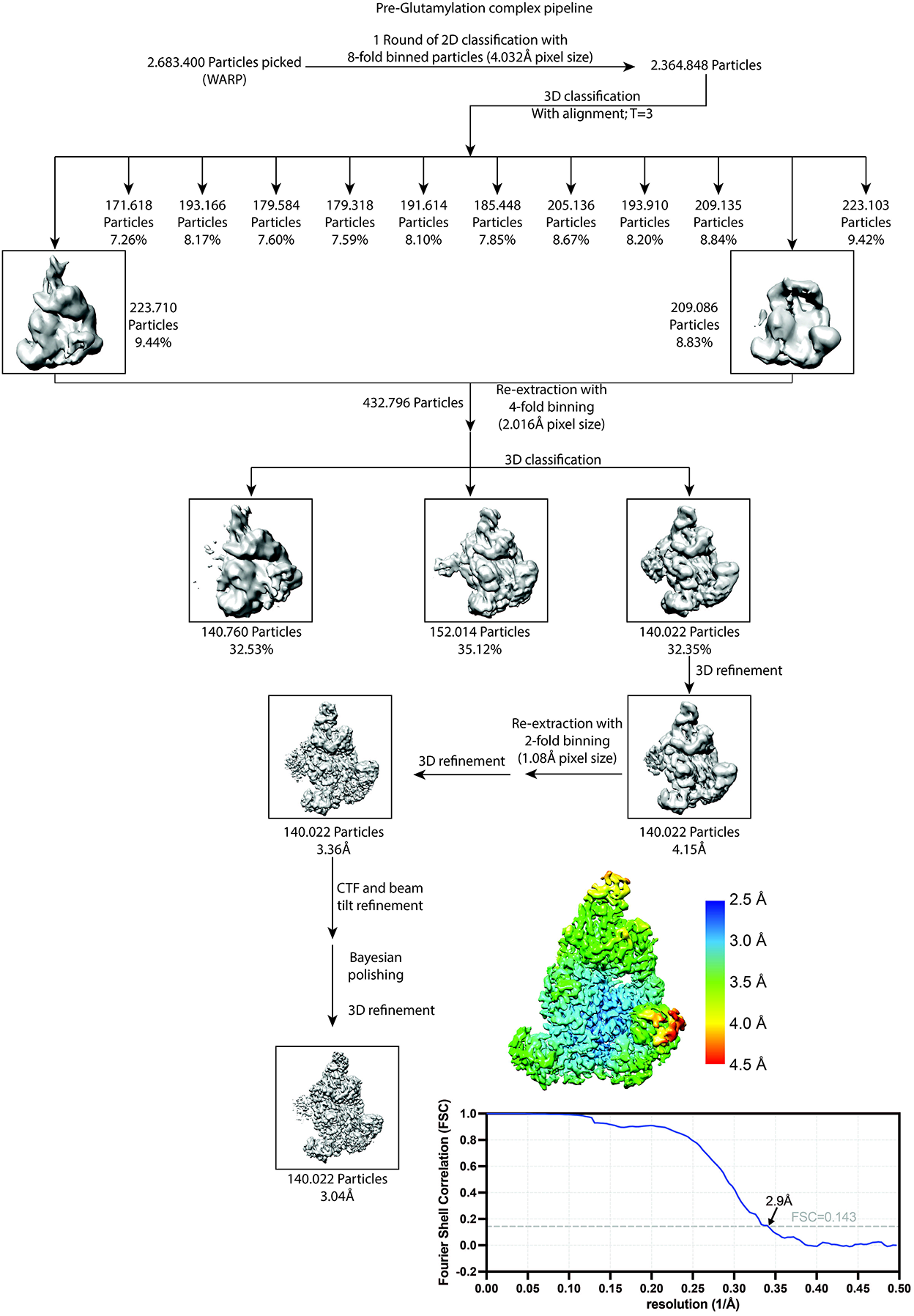
Processing pipeline of SidJE-565A/CaM/SdeA reaction intermediate complex Particles were picked using WARP and processed using RELION, pipeline shows models up to final 3D refinement step. Bottom right, Gold Standard FSC curves of SidJ-E565A/CaM/SdeA pre-glutamylation complex, and final map of SidJE-565A/CaM/SdeA colored according to the local resolution estimates.

**Figure S3.**
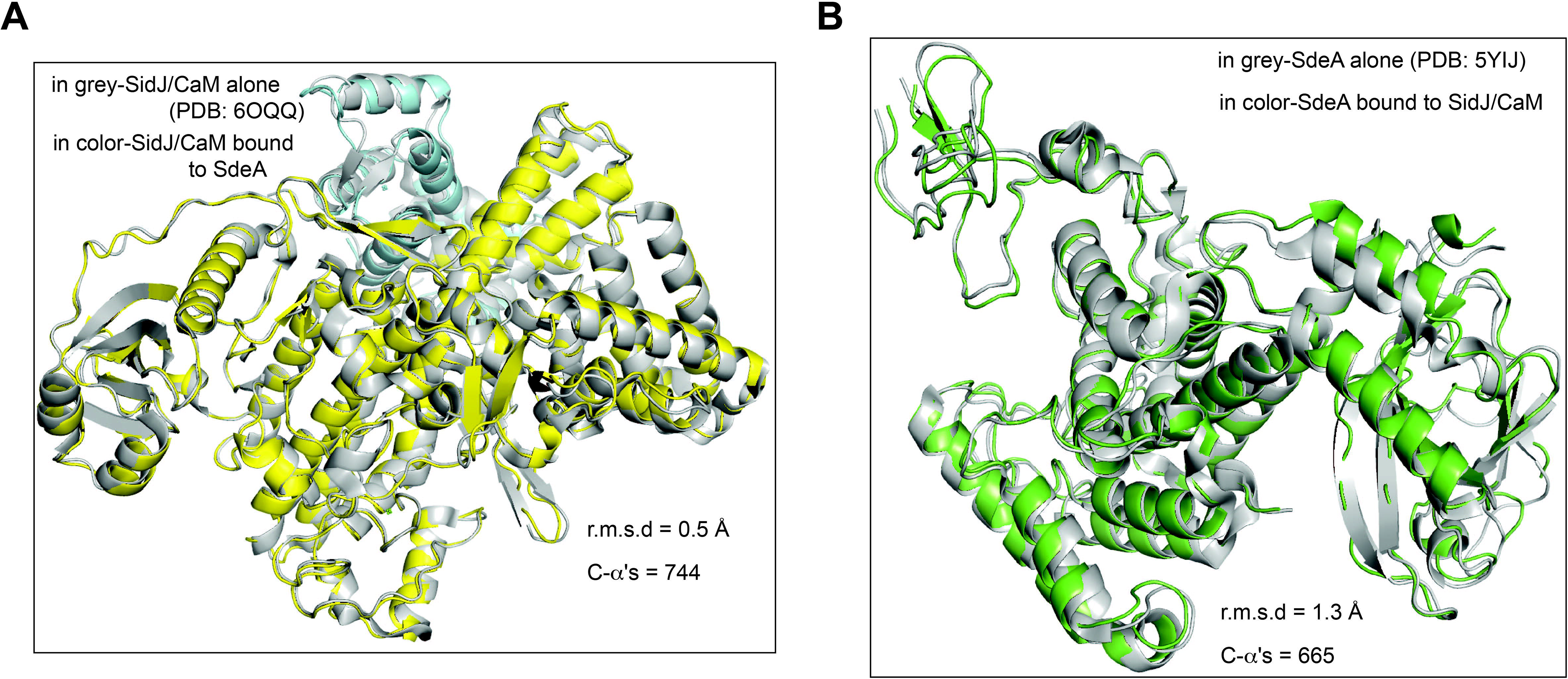
Structural comparison of SidJ and SdeA alone and in complex A) Comparison between SidJ/CaM crystal structure (Grey, PDB: 6OQQ) and SidJ/CaM in the pre-glutamylaiton complex SidJ-E565A/CaM/SdeA (Yellow, Teal) B) Comparison between SdeA crystal structure (Grey, PDB: 5YIJ) and SdeA in the the pre-glutamylaiton complex SidJ-E565A/CaM/SdeA (Green)

**Figure S4.**
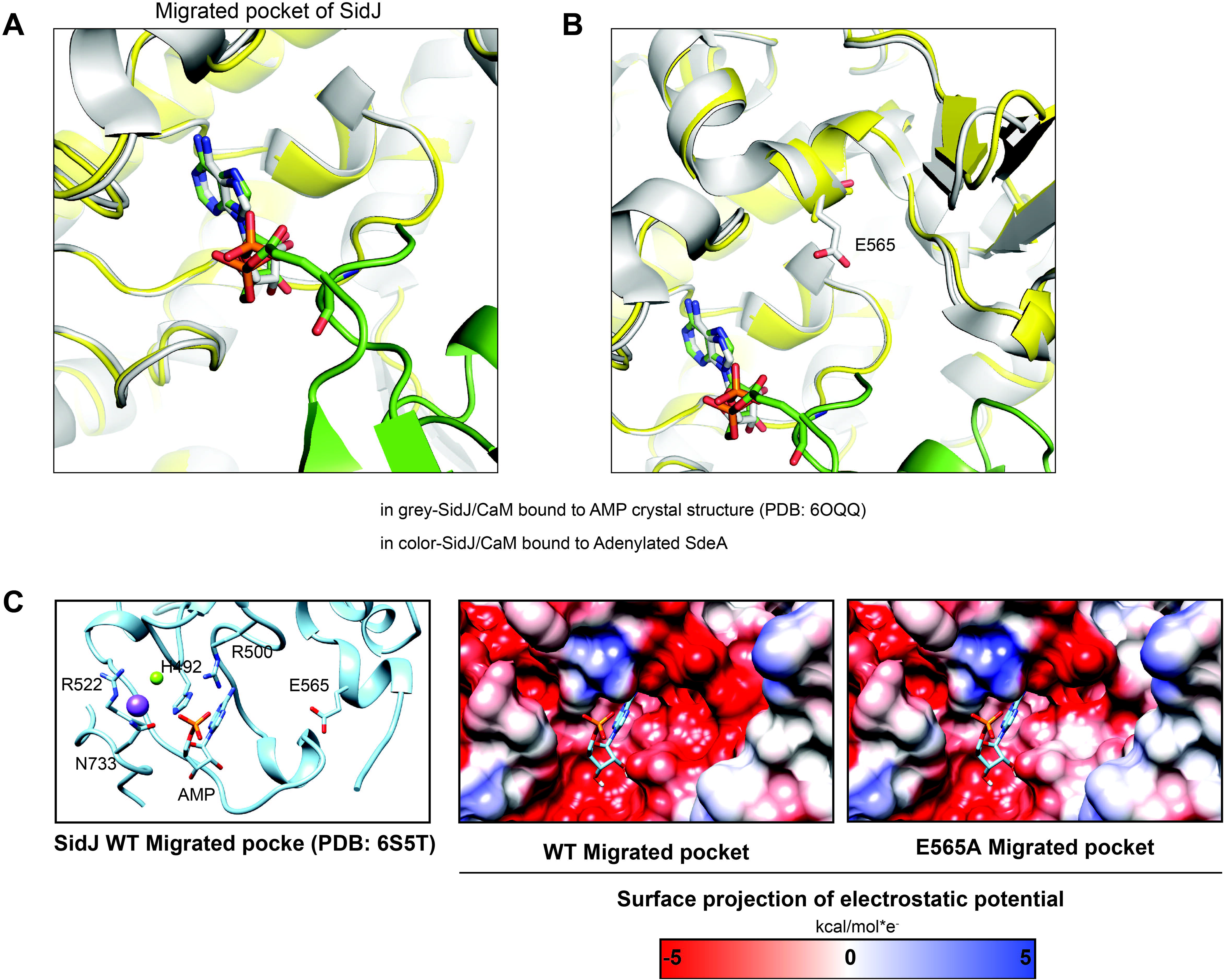
SidJ mutation E565A does not induce structural change A) Comparison of SidJ/CaM migrated pocket with AMP bound in crystal structure (Grey, PDB: 6OQQ) and SidJ-E565A/CaM/SdeA with AMPylated E860 bound in the catalytic intermediate complex (Colored) B) Comparison of SidJ/CaM migrated pocket structure in WT-SidJ crystal structure (Grey) and SidJ-E565A/CaM/SdeA catalytic intermediate complex C) Electrostatic potential is projected onto the surface of SidJ WT (PDB: 6S5T) and SidJ E565A (modeled) migrated pocket. AMP is modeled based on the crystal structure of SidJ (PDB: 6OQQ)

**Figure S5.**
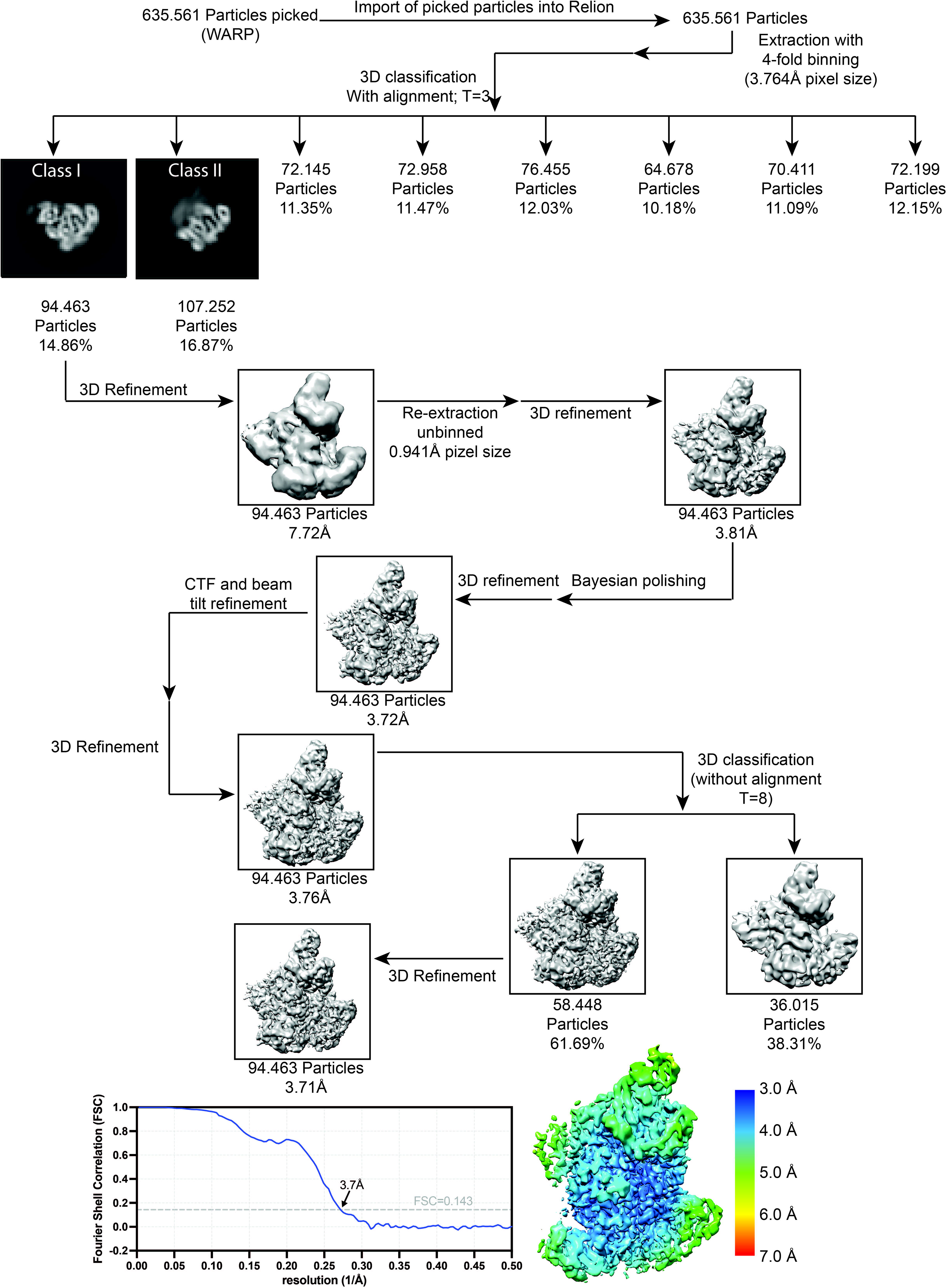
Processing pipeline of SidJE-565A/CaM/SdeA post-catalytic complex Particles were picked using WARP and processed using RELION, pipeline shows models up to final 3D refinement step. Bottom left, Gold Standard FSC curves of SidJ-E565A/CaM/SdeA post-catalytic complex. Bottom right, Final map of the post-catalytic SidJE-565A/CaM/SdeA complex colored according to the local resolution estimates.

**Figure S6.**
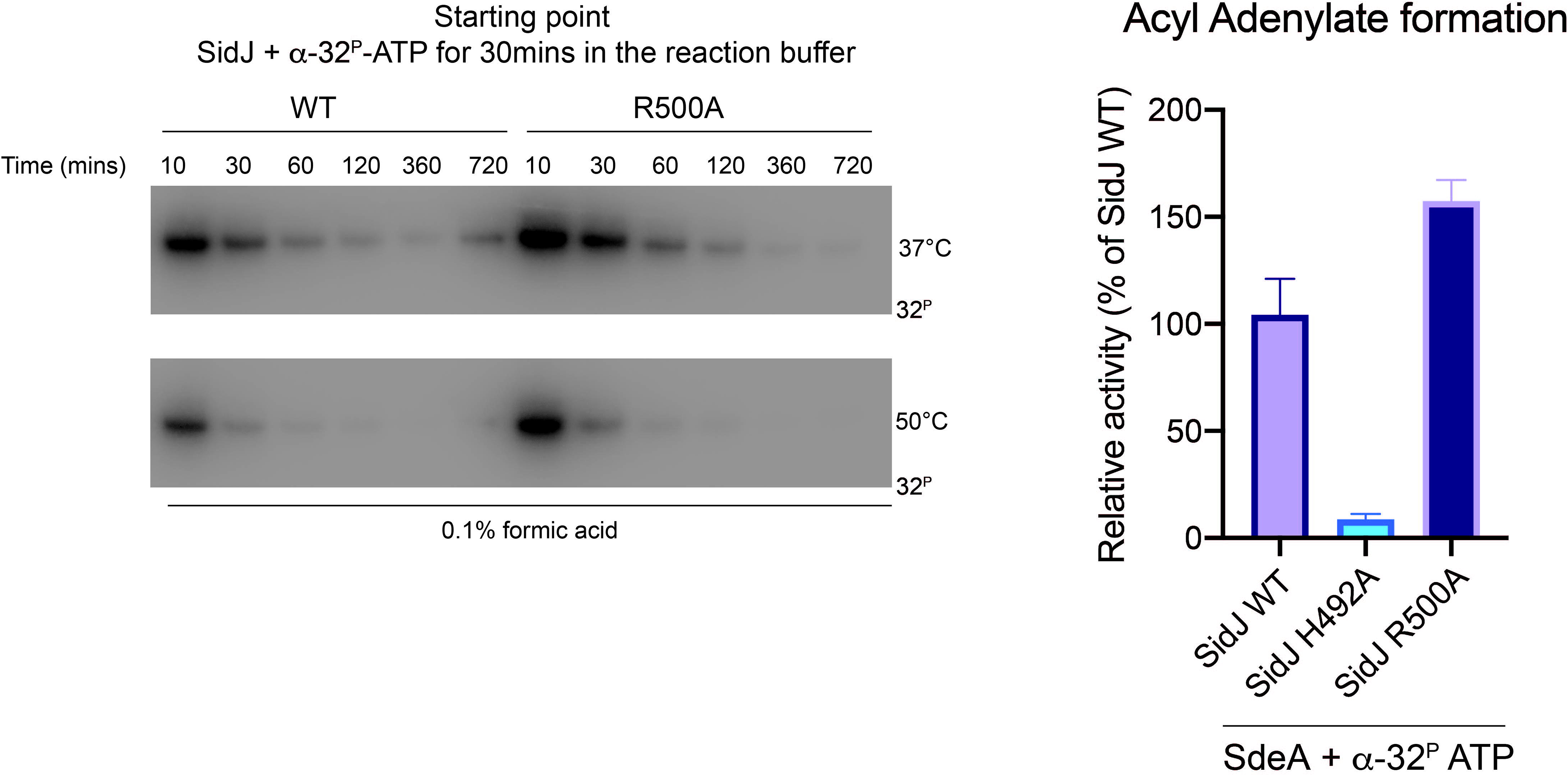
Characterization of SidJ R500A mutant Left, incorporation of α-[^32^P]-ATP into SidJ or SidJ R500A. After the reaction the samples were treated at the indicated temperature for various time points in 0.1% formic acid. Reactions were separated by SDS-PAGE and visualized by autoradiography. Data shown in figure 5b is a cropped version of data shown in this figure. Right, Acyl adenylylate formation after reaction with SidJ WT or mutants in the presence of WT SdeA and α-[32P]-ATP. Reactions were terminated by the addition of TCA, and TCA-insoluble pellets were measured by scintillation counting.

**Figure S7:**
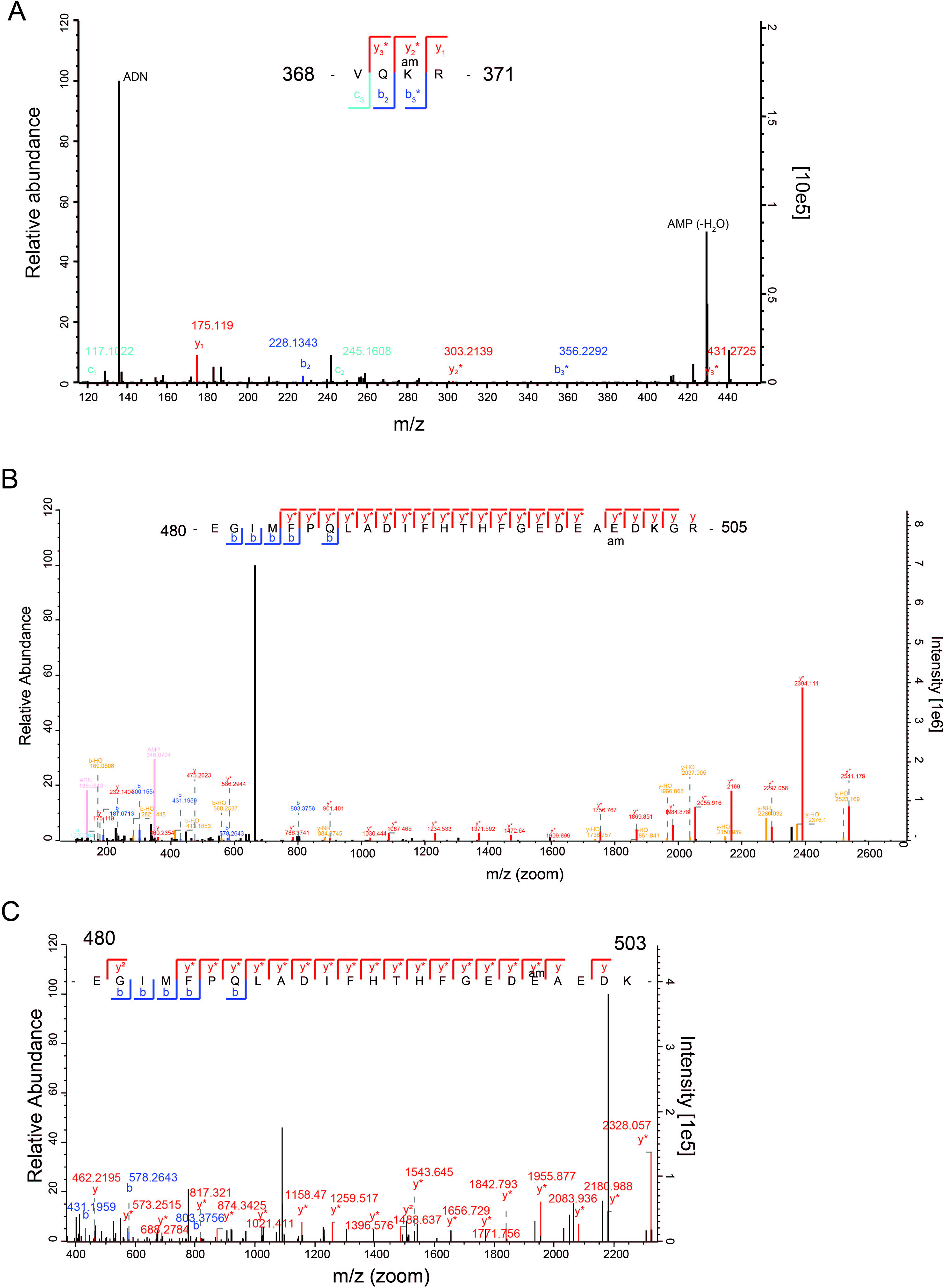
Mass spectrometry analysis of SidJ-autoAMPylation. A) LC-MS/MS of SidJ R500A autoAMPylation reaction showing the HCD fragmentation spectra of the canonical pocket peptide. B) LC-MS/MS of SidJ R500A autoAMPylation reaction showing the HCD fragmentation of the bridging peptide. C) HCD Fragmentation spectrum of SidJ R500A peptide E480-K503 with a possible AMPylation site E499 highlighted

**Figure S8.**
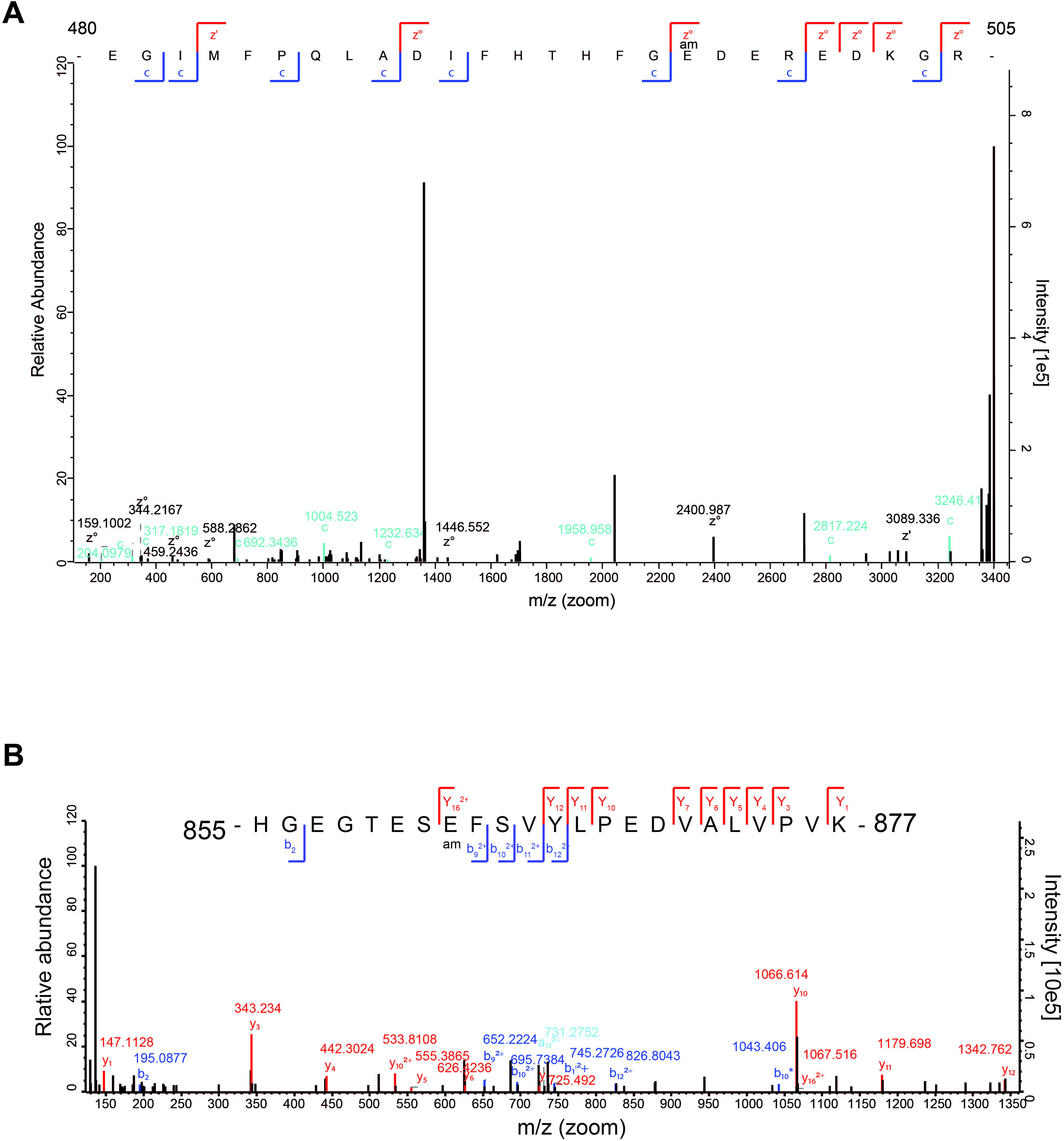
HCD fragmentation spectra showing the AMPylation of SdeA mART catalytic peptide. A) LC-MS/MS of SidJ WT autoAMPylation reaction showing the ETD Fragmentation of the bridging peptide. B) HCD fragmentation spectrum of SdeA catalytic peptide. A reaction was set up with SidJ R500A/CaM with SdeA in the presence of ATP. The reaction components were subjected to LC-MS/MS.

**Figure S9.**
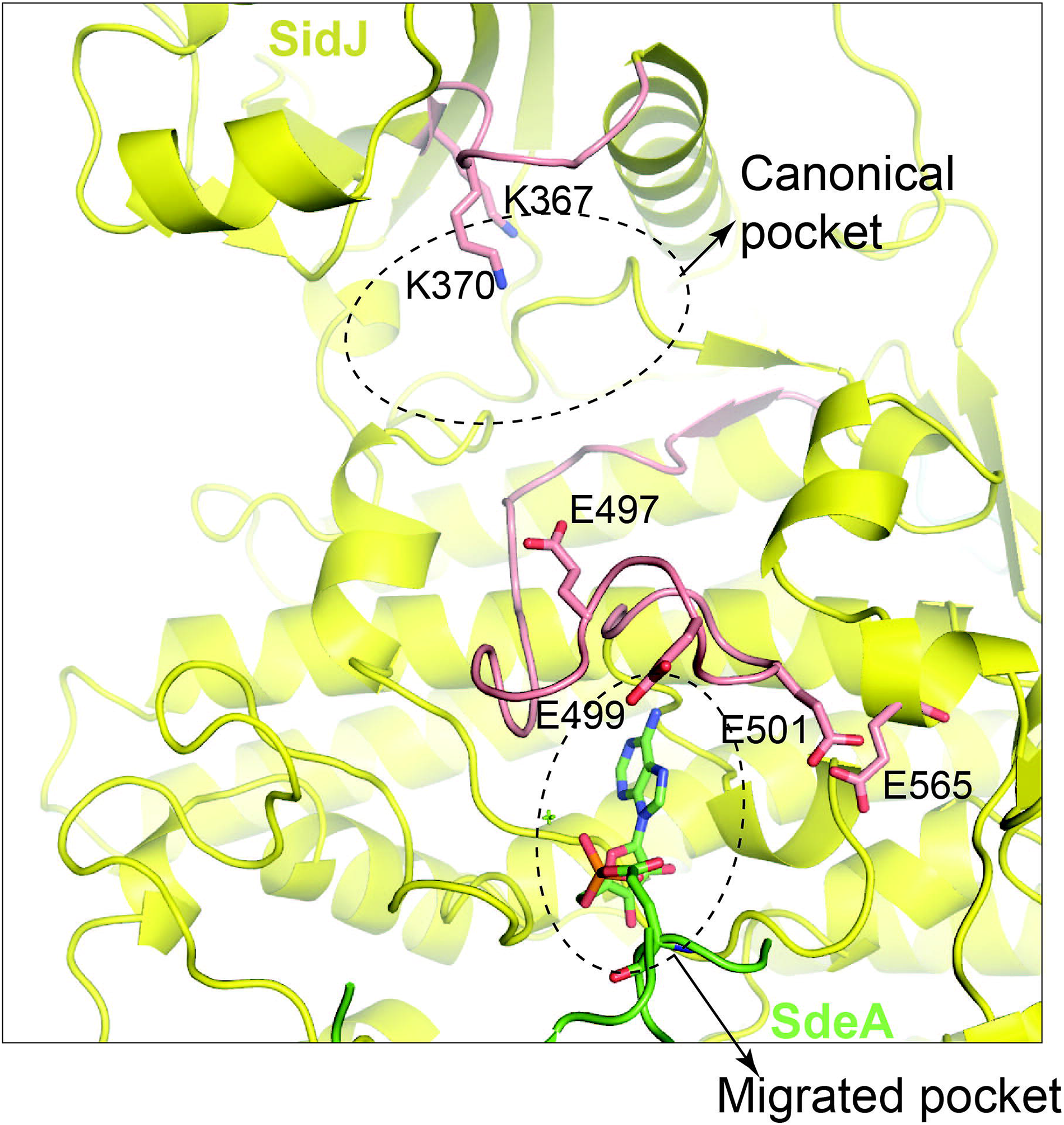
SidJ structure showing the cluster of glutamate residues around the autoAMPylation sites found by mass spectrometry. Detailed view of SidJ auto-AMPylation sites, with peptides recovered in mass spec shown in red, modification sites including a series of glutamates extending into the migrated pocket are highlighted

**Figure S10.**
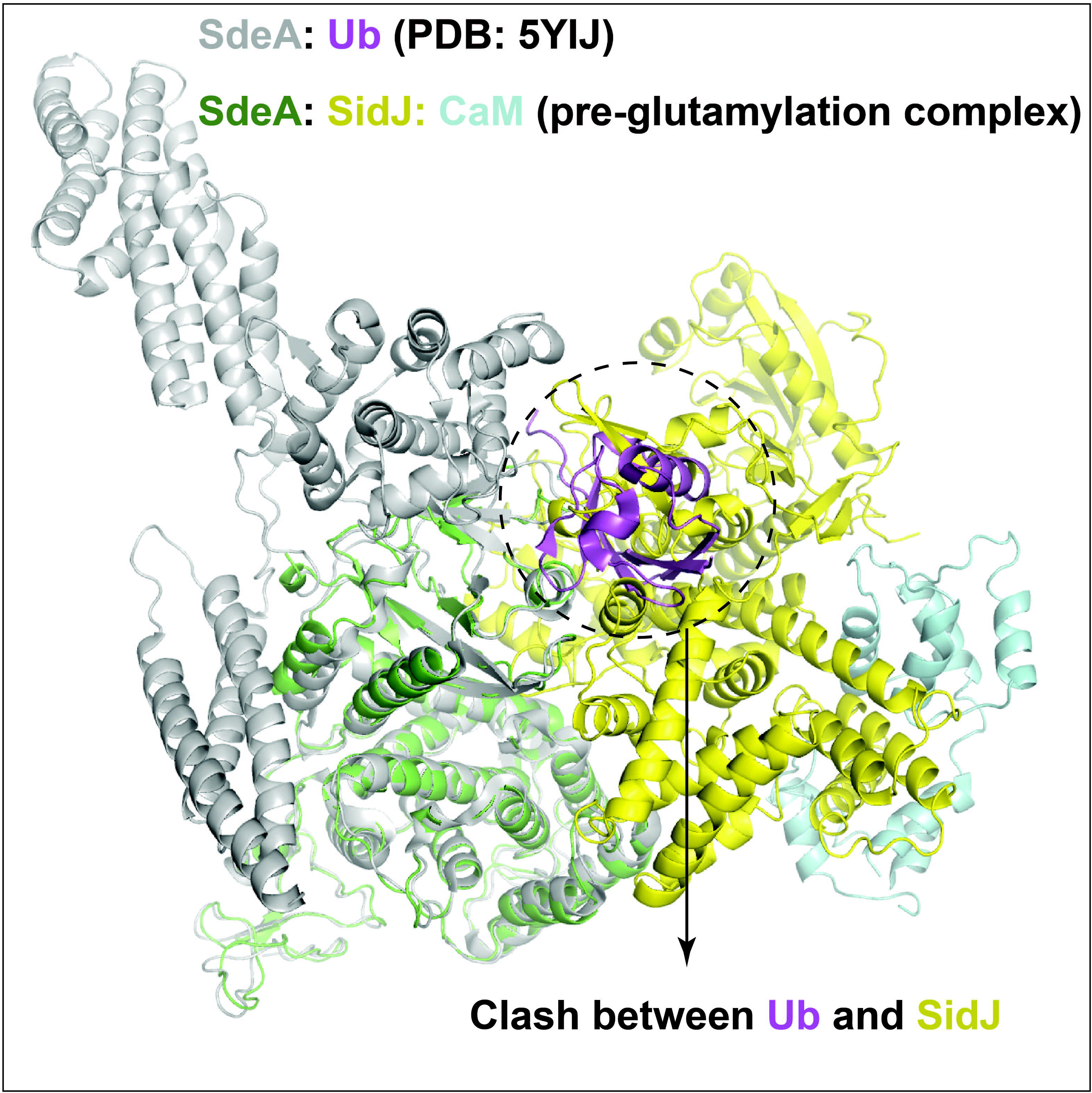
SidJ and Ubiquitin compete for the same binding surface on the mART domain of SdeA. SdeA-Ub complex structure (PDB: 5YIJ) is superposed on the pre-glutamylation complex structure of SidJ/CaM-SdeA.

## Materials and methods

### Purification of SidJ/CaM and SdeA proteins

Expression and purification of SidJ/CaM and SdeA constructs were performed using the protocol published in Bhogaraju et al 2019^16^. Briefly, constructs of SidJ 99-C or SdeA 231-1190 were cloned into pCoofy1 vector, while FL CaM was cloned into pET15c. The validated constructs were transformed into chemically competent BL21 Star (Sigma Aldrich) using heat-shock method, with SidJ being co-expressed with Calmodulin, and SdeA being expressed alone. The cells were grown LB at 37°C until OD600=0.6, induced using 0.5mM IPTG, and expressed for 18 hours at 18°C and harvested. The resulting pellets were re-suspended in Lysis buffer (300mM NaCl, 50mM Tris pH 7.5, 10% Glycerol) containing the protease inhibitor cocktail (Roche), and lysed using sonication. The lysed cells were then centrifuged at 13,000RPM, filtered using a 0.22 μm filter, and applied to 3mL of Talon bead resin (Takara) that was equilibrated into lysis buffer. The clarified lysate was incubated under gentle agitation at 4°C for 60 minutes, and centrifuged at 500g for 2 minutes. The supernatant was decanted, and the beads were washed and incubated for 10 minutes with lysis buffer three times, each time centrifuging and decanting the supernatant. Further impurities were removed through the addition and incubation of Lysis buffer with 10mM Imidazole. The protein was then eluted in multiple steps using imidazole concentrations of 50mM to 300mM, and purity was determined using SDS-PAGE. The pure fractions were concentrated, and loaded onto a pre-equilibrated (100mM NaCl, 10mM HEPES pH 7.5, 0.5mM TCEP) Superdex S200 Increase 10/300 size exclusion column (GE Life Sciences). Peak fractions were evaluated via SDS-PAGE and the purest fractions were pooled, concentrated to 1.5mg/mL and aliqots flash-frozen.

### Preparation of trapped SidJ/CaM-SdeA heterotrimer for cryo-EM (GRAFIX)

The sample was trapped and cross-linked using the GraFix protocol^22^. Equimolar quantities of SidJ-E565A/CaM and SdeA were incubated on ice for 30 minutes in the presence of 150mM NaCl, 50mM HEPES, 10mM MgCl2, and 10mM ATP and loaded onto 5-20% Glycerol gradients using 0%-0.1% Glutaraldehyde and 100mM NaCl, 10mM HEPES pH 7.5, 10mM MgCl2, and the tubes were centrifuged at 20.000rpm for 18 hours at 4 °C. The gradients were subsequently fractionated and quenched through the addition of 10mM Tris (final concentration). The fractions were evaluated by SDS-PAGE and silver staining using SilverQuest Silver Stain Kit (Invitrogen). The fractions containing a band of the desired weight were pooled, concentrated, and loaded onto a pre-equilibrated (150mM NaCl, 10mM HEPES pH 7.5, 10mM MgCl2, 0.5mM TCEP) Superdex S200 3.2/300 size exclusion column (GE Life Sciences). The peak fractions were pooled and concentrated to 0.3-0.5 mg/mL and used for grid preparation.

### Vitrification and Cryo-EM

2μL of the cross-linked and concentrated sample was applied using a Vitrobot MkIV (Thermo Scientific) to each side of a Quantifoil Au 300 1.2/1.3 grid, which was glow discharged using a Pelco EasyGlow on both sides using 30mA for 30 seconds before being blotted using 100% humidity at 4 °C and blot force −4 or −6 for 2 seconds and being plunge frozen in liquid ethane. In the case of the post-catalytic structure, 5mM of Glutamate (final concentration) were added to the sample immediately prior to application to the grid.

The grids were screened using a Glacios 200kV microscope (Thermo Fisher) with a Falcon III direct electron detector (Thermo Fisher). Movies for the post-catalytic complex were collected in electron counting mode (defocus range of −1.0μm to −2.5μm, 0.25μm step size) with a magnified pixel size of 0.941 Å at a dose rate of 0.93 e/Å2/s for a total dose of 35 e/Å2 fractioned over 30 movie frames. Using the same parameters, a short data collection was performed on a grid containing the pre-glutamylation complex, in order to generate an ab initio model for data processing.

For the pre-glutamylation SidJ/CaM-SdeA complex, the movies were collected on a Titan Krios (FEI) equipped with a Quantum-K3 detector (Gatan), a defocus between −0.7μm and −1.7μm was applied in 0.1μm steps during collection in electron counting mode. A magnified pixel size of 0.504 Å, and a dose rate of 15 e/pix/s for a total dose of 47.45 e/Å2 were used, distributed over 40 frames.

### Cryo-EM data processing

To generate an initial model for the catalytic intermediate complex, movies collected on the Glacios 200 kV microscope (Thermo Fisher) were used for particle picking in WARP using the BoxNet2Mask_20180918 model, and imported into Relion 3.1^32^ for ab initio 3D-classification. The best class was chosen and scaled to serve as the reference model for the high-resolution structure.

For the pre-glutamylation complex, motion correction and contrast-transfer function (CTF) estimation, with subsequent particle picking using the BoxNet2Mask_20180918 model were performed in WARP^33^. Coordinates of 2,683,400 particles were imported into Relion 3.1 and initially extracted with an 8-fold binning factor. After one round of reference-free 2D classification, 2,364,848 particles were included for 3D-classification, with the previous low-resolution map as a reference. A subset of 432.796 particles from the best two classes were re-extracted with a binning factor of 4, classified again, and 140,022 particles from the best class were picked and refined to 4.15 Å resolution. The particles were once again re-extracted using a binning factor of 2, and refined to a resolution of 3.36 Å, before CTF refinement, beam-tilt correction, and Bayesian particle polishing were performed. These steps resulted in a model resolution of 2.94 Å. The reported overall resolution of 2.9 Å was calculated using the gold-standard Fourier shell correlation (FSC) 0.143 criterion^34^ and was corrected for the effects of a soft mask on the FSC curve using high-resolution noise substitution^35^.

For the post-catalytic complex, motion correction and contrast-transfer function (CTF) estimation, with subsequent particle picking using the BoxNet2Mask_20180918 model were performed in WARP. Coordinates of 635,561 particles were imported into Relion 3.1 and initially extracted with an 4-fold binning factor. All imported particles were included for 3D-classification to generate an ab initio model. A subset of 94,463 particles from the best class were refined to 7.72 Å resolution. The particles were re-extracted without binning, and refined to a resolution of 3.81 Å, before CTF refinement, beam-tilt correction, and Bayesian particle polishing were performed. After another round of 3D-refinement, which produced a model of 3.76 Å resolution, the particles were classified one more time, and 58,448 particles from the best class were picked for a final round of 3D-refinement. These steps resulted in a map resolution of 3.71 Å.

### Immunoprecipitation

HEK293 cells were co-transfected transiently with mCherry-SidJ (pmCherry-C1 vector) and either GFP (pEGFP-C1 vector) or GFP-SdeA (pEGFP-C1) using polyethylenimine. Cells were harvested at 18 hours post-transfection, washed with PBS (phosphate buffered saline) and lysed in immunoprecipitation buffer (50 mM Tris-HCl pH 7.5, 150mM NaCl, 1% Triton X-100, and protease inhibitor cocktail (cOmplete Mini EDTA-free from Roche) and mixed with GFP-Trap Agarose beads (ChromoTek) and incubated for 2 hours at 4°C while being subjected to end-to-end rotation. The beads were washed three times with the immunoprecipitation buffer. Finally, proteins were eluted by boiling with 4x Laemmli buffer for 10 minutes, then separated through SDS-PAGE, and visualized following western blot. The Antibody used for mCherry detection was DSRed2 sc-101256 (Santa Cruz Biotechnology), and GFP was detected with GFP sc-9996 (Santa Cruz Biotechnology).

### Glutamylation assay

20uL in vitro glutamylation reactions were performed with 2uM SidJ/CaM and 2uM SdeA, 5mM ATP and 50uM (0.5 nCi) L-[14C]Glutamate (Perkin Elmer) in a buffer consisting of 30mM HEPES pH 7.5, 150mM NaCl, 10mM MgCl2, 1mM β-Mercaptoethanol and incubated at 37 °C for 30 minutes. Reactions were stopped through the addition of 5uL of 4x SDS-PAGE sample buffer. The samples were then separated using SDS-PAGE, stained using Coomassie stain, and dried using a Model 583 gel dryer (Bio-Rad). A storage phosphor screen (GE Healthcare) was placed on the gel and exposed for 72 hours, and the autoradiography signal was collected using a Typhoon FLA 7000 (GE).

### Acyl Adenylate formation assay

AMPylation was measured using the protocol described in Black et al (2019). Briefly, reactions were carried out in the presence of 150μM (5μCi) [α-32P]ATP. The 20μL reactions contained 1mg/ml bovine serum albumin (BSA), 100mM Sodium Acetate, 150mM NaCl, 50mM Tris, 50mM Bis-Tris pH 6.5, 0.5 mM MgCl2, and 1mM DTT, and equimolar quantities of 5μM SidJ/CaM and SdeA. Reactions were carried out on ice, and stopped after 30 minutes by adding 500μL of ice-cold 20% TCA, and incubated on ice for 40 minutes. Products were centrifuged at 21,000g for 15 minutes, before washing the pellet twice with 250μL of ice-cold TCA. The radioactivity of the acid-insoluble pellet was measured through the addition of 50μL of MicroScint PS (Perkin Elmer), vortexing, and performing scintillation counting on a MicroBeta 2450 Microplate Counter (Perkin Elmer). The graphs were prepared using GraphPad Prism.

### Auto-AMPylation assay

20μL in vitro glutamylation reactions were performed with 2μM SidJ/CaM, 2.5μCi α - [32P]ATP (Perkin Elmer) in a buffer consisting of 30mM HEPES pH 7.5, 150mM NaCl, 10mM MgCl2, 1mM β-Mercaptoethanol and incubated at 37 °C for 30 minutes. Reactions were stopped through the addition of 5μL of 4x SDS-PAGE sample buffer. The samples were then separated using SDS-PAGE, stained using Coomassie stain, and dried using a Model 583 gel dryer (Bio-Rad). A storage phosphor screen (GE Healthcare) was placed on the gel and exposed for 18 hours, and the autoradiography signal was collected using a Typhoon FLA 7000 (GE).

### Analytical gel filtration

Analytical gel filtrations were performed using equimolar mixtures of SidJ/CaM constructs and SdeA constructs at a concentration of 0.5mg/mL. Under the addition of 10mM MgCl and 5mM ATP, the sample was incubated on ice for 30 minutes before being loaded onto a pre-equilibrated (100mM NaCl, 10mM HEPES pH 7.5, 100mM NaCl 10mM MgCl2, 0.5mM TCEP) Superdex S200 3.2/300 size exclusion column (GE Life Sciences). The resulting chromatograms were overlaid and compared to calibration curves to estimate molecular weight. Chromatograms were plotted using GraphPad Prism.

### Mass spectrometry

Reaction mixtures were digested through the addition of 500ng of Promega Trypsin Gold (resuspended in 500nl of 50mM Acetic acid) and 20 ul 50mM ammonium bicarbonate (ABC). After 90 minutes incubation at room temperature digests were applied to in-house manufactured C18 StageTips equilibrated with 10mM ABC. Samples were washed with 10mM ABC, eluted with 40% acetonitrile in 10mM ABC and dried for 1 hr by vacuum centrifugation.

LC-MS analyses were performed using either an Easy-nLC 1000 coupled to an Orbitrap Fusion mass spectrometer, or an Easy-nLC 1200 coupled to an Orbitrap Fusion Lumos (Thermo Scientific) with peptides generated from roughly 500ng of proteins injected for each analysis onto either 75 micron x 25cm packed emitter columns (New Objective) or 75 micron x 50 cm C18 Acclaim Pepmap columns (Thermo Scientific). Peptides were eluted with linear gradients from 1% to 35% solvent B (80% Acetonitrile in 0.1% formic acid) in either 30 or 45 minutes, followed by a steeper wash phase.

In general, samples were analyzed with “Topspeed” data-directed analysis methods with a cycle time setting of 2 or 3 seconds. In order to obtain high-quality fragmentation data, the MS2 AGC fill was set high for both HCD (300-500%) and ETD (200%) acquisitions. In some acquisitions rapid HCD fragmentation was performed with standard parameters to screen for AMPylated peptides. The presence of an Adenine (ADN) fragment ion (136.062 Da) was used to trigger higher-quality HCD or ETD fragmentation.

Due to the possible lability of the AMP modification, various acquisition strategies were applied. Higher-energy collision induced decay (HCD) was applied initially in order to identify modified peptides. This mode of fragmentation tends to break off the modifier. Interestingly two fragmentation behaviors were observable. Diagnostic ions for breakage of AMP appeared at either 330.060 (representing the broken-off AMP modifier itself in the H+ state) or 348.071 (AMP having taken an additional H2O from the sidechain). In the peptides modified on Lysine, the AMP+H2O ion does not appear, presumably since there is no oxygen involved in binding to the Lysine sidechain. In those peptides where the modifier appears to be on E, the heavier diagnostic ion is observed. We would propose that this heavier diagnostic ion is characteristic of O-linked AMPylation.

### Pyrophosphate release assay

The ATPase activity of WT SidJ and its mutants was measured in a UV-transparent microplate using the EnzChek Pyrophosphate Assay Kit (ThermoFisher Scientific, E-6645). The assay was performed in triplicates. All the components were added in the reaction mixture as described in the kit. 5μM of WT SidJ, and mutants were added into the reaction mixture. Finally, 2 mM ATP was then added into the reaction mixture to start the reaction and absorbance measurements at 360 nm were taken immediately and continuously at 1-min intervals using a Clariostar plate reader.

## Acknowledgements

We thank Sarah Gharbi for technical assistance. We thank Ivan Dikic, Donghyuk Shin, Sissy Kalayil, Simonne Griffith-Jones, Mohit Misra and Wojtek Galej for the critical reading of the manuscript. We thank Matthew Bowler for help with generating the AMPylated glutamate residue cif file. We thank the Partnership for Structural Biology (PSB) biophysical platform, especially Philippe Mas and Caroline Mas for the training and support. We thank the EMBL Mass spectrometry core facility, especially Mandy Rettel and Frank Stein for the data acquisition and analysis. We thank Alice Aubert and Martin Pelosse for training and maintenance of the EMBL eukaryotic expression facility. We thank Sarah Schneider, Erika Pellegrini and Michael Hons for training and maintenance of EMBL Grenoble EM facility.

## Author Contributions

M.A. performed protein purification, cryo-EM, radioactive experiments. R.S. performed protein purification, biochemistry. T.C. performed the mass spectrometry experiments under the supervision of I.M. F.W. collected the cryo-EM data for pre-glutamylation structure. S.B. supervised the study and wrote the paper with inputs from all the authors.

---

**Table S1:**
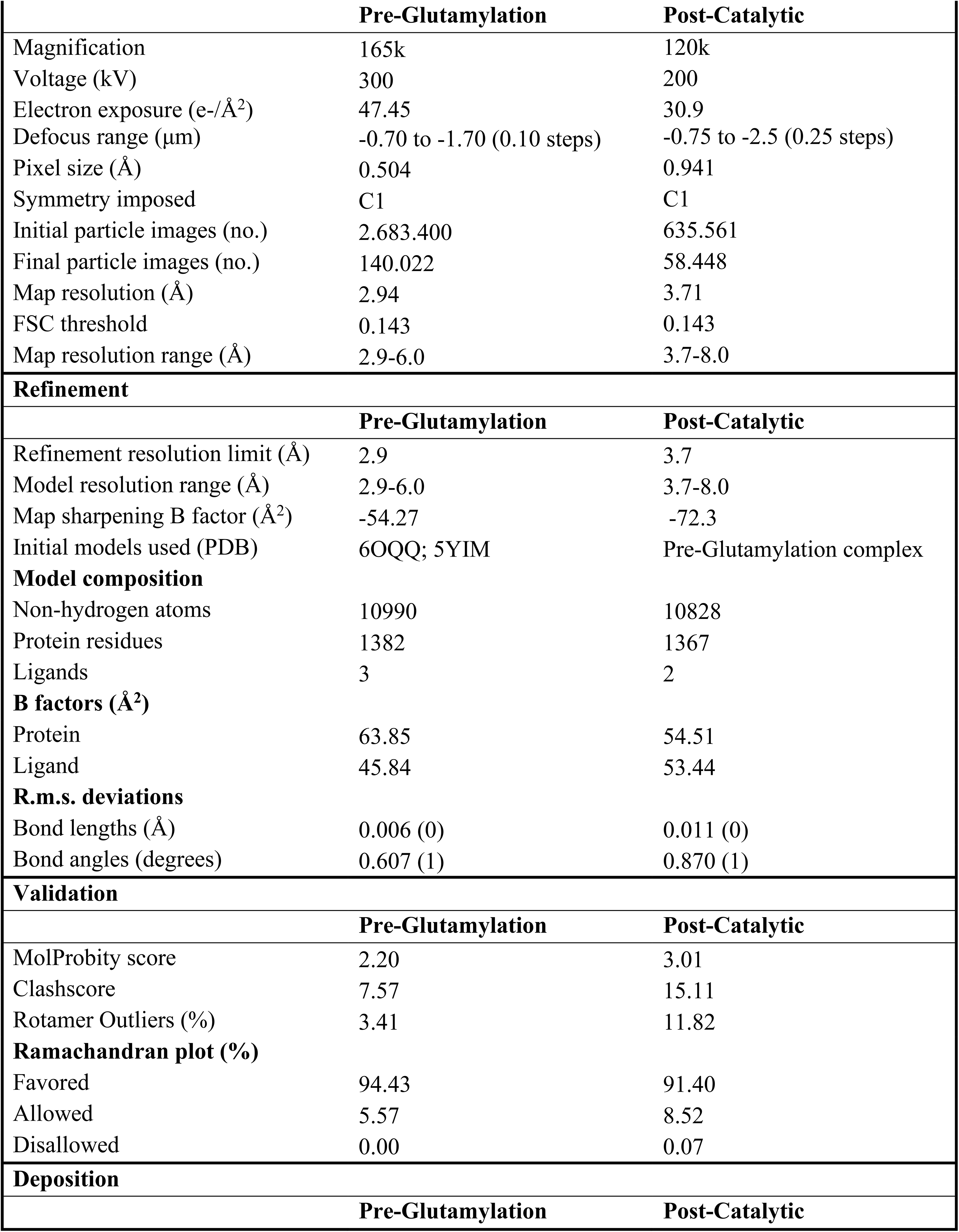
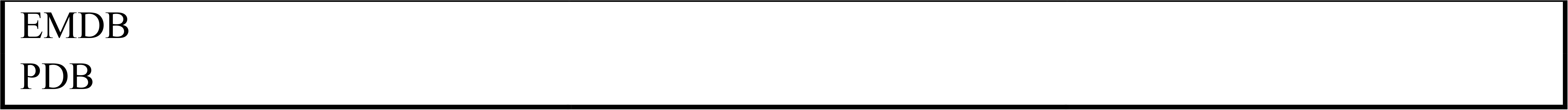
cryo-EM data collection and processing parameters.

